# The protective effect of aspirin in colorectal carcinogenesis: a multiscale computational study from mutant evolution to age incidence curves

**DOI:** 10.1101/2021.05.11.443671

**Authors:** Yifan Wang, C. Richard Boland, Ajay Goel, Dominik Wodarz, Natalia L. Komarova

## Abstract

Aspirin intake has been shown to lead to significant protection against colorectal cancer, e.g. with an up to two-fold reduction in colorectal adenoma incidence rates at higher doses. The mechanisms contributing to protection are not yet fully understood. While aspirin is an anti-inflammatory drug and can thus influence the tumor microenvironment, in vitro and in vivo experiments have recently shown that aspirin can also have a direct effect on cellular kinetics and fitness. It reduces the rate of tumor cell division and increases the rate of cell death. The question arises whether such changes in cellular fitness are sufficient to significantly contribute to the epidemiologically observed protection. To investigate this, we constructed a class of mathematical models of in vivo evolution of advanced adenomas, parameterized it with available estimates, and calculated population level incidence. Fitting the predictions to age incidence data revealed that only a model that included colonic crypt competition can account for the observed age-incidence curve. This model was then used to predict modified incidence patterns if cellular kinetics were altered as a result of aspirin treatment. We found that changes in cellular fitness that were within the experimentally observed ranges could reduce advanced adenoma incidence by a sufficient amount to account for age incidence data in aspirin-treated patient cohorts. While the mechanisms that contribute to the protective effect of aspirin are likely complex and multi-factorial, our study demonstrates that direct aspirin-induced changes of tumor cell fitness can significantly contribute to epidemiologically observed reduced incidence patterns.

## Introduction

Colorectal cancer currently affects about 5% of the population in the USA and is a major cause of cancer-related deaths [1]. Prevention of colorectal cancer is an important goal in the quest to reduce morbidity and mortality. In this respect, long-term aspirin use has been shown to be effective [2,3]. Aspirin is a non-steroidal anti-inflammatory drug (NSAID) and is a cyclo-oxygenase (COX)-2 inhibitor [4]. The CAPP2 trial [5] demonstrated that the intake of 600mg of aspirin per day for 2 years resulted in a 63% reduction in colorectal cancer incidence in Lynch Syndrome patients. Interestingly, observation of the protective effect of aspirin required a follow-up time of more than 55 months [5]. The mechanisms underlying this protective effect therefore are probably complex and multi-factorial. Inflammation is a likely driver of colorectal carcinogenesis [6], and aspirin can reduce the extent of inflammation in the cellular microenvironment, which might contribute to a reduced development of disease. Our previous in vitro and in vivo work, however, has shown that physiologically relevant aspirin concentrations can also have a direct effect on tumor cells, reducing their rate of proliferation and increasing their death rate [7,8]. This not only results in reduced tumor growth, but can also lead to a lower probability that newly generated tumor cells successfully give rise to clonal expansion, thus increasing the likelihood that these initially transformed cells go extinct [9]. This effect might contribute to the reduced incidence of colorectal cancer as a result of aspirin intake.

While these direct effects of aspirin on tumor cell division and death rates have been documented in vitro and in vivo [7,8], and occurred under physiologically realistic doses, it is unclear to what extent these changes in cellular kinetics can potentially alter disease incidence. To evaluate this quantitatively, a mathematical modeling framework needs to be developed that predicts epidemiological incidence data based on cellular processes. There is a rich history of such approaches in the cancer literature in different contexts [10,11,12,13,14,15,16], which has allowed researchers to gain fundamental insights into carcinogenic processes based on the interpretation of age-incidence data. Here, we describe a mathematical model of advanced adenoma formation and parameterize it by fitting epidemiological predictions to incidence data that document advanced adenoma occurrence as a function of age. We then use this model to test whether aspirin-mediated changes in cellular kinetics, as documented by our experiments, can result in reductions in advanced adenoma incidence that are comparable to those observed in aspirin-treated patient cohorts. We find that the magnitude of changes in the kinetics of transformed cell populations that we observed experimentally can result in a pronounced reduction of advanced adenoma incidence, and that the epidemiologically observed incidence reductions (up to 50%) can be explained by our model. This indicates that the direct effects of aspirin on dividing cells can in principle explain much of the chemoprotective effect exerted by this drug. We note, however, that while this is a clear result that emerges from this mathematical modeling effort, other mechanisms of aspirin not included in this model (such as anti-inflammatory effects) are likely to also contribute to the observed protective effect.

We start by describing a mathematical model of advanced adenoma formation and show that when parameterized with experimentally obtained estimates, it can account for epidemiologically observed age-incidence curves, only as long as inter-crypt competition is explicitly included. We then use this model to simulate the effect of aspirin on the incidence of advanced adenomas in human populations, and compare model predictions to epidemiological data.

## Methods

### Computational modeling

In order to quantify the effects of aspirin on colorectal cancer initiation and progression, we have designed a mathematical model that is rooted in the process of multistep carcinogenesis [13,16,17,18]. Its assumptions are similar in principle to those in a recent study [19], with important differences that are discussed below. There are two early molecular events that we postulate (without assuming their temporal order): (1) An inactivation of the APC gene, or a related event that affects the functioning of the beta-catenin/WNT signaling pathway, and (2) an activation of the KRAS oncogene. The inactivation of the APC tumor suppressor gene is a classic example of a loss-of-function mutation, which implies two molecular events, corresponding to the inactivation of the two copies of the gene. The associated mutation rate is therefore assumed to be u=10^−7^ per cell division. The activation of the KRAS oncogene on the other hand is a gain-of-function event, whose mutation rate is about two orders of magnitude lower (μ=10^−9^ per cell division). The associated selection-mutation diagram is shown in **Fig 1** and contains six different cell populations, denoted as types 1 through 6. The populations occupying the top row (types 1--3) are characterized by an unmutated KRAS oncogene; the populations of the bottom row (types 4-6) all have the KRAS mutation activated. Moving from left to right on this diagram, the number of inactivated copies of the APC gene increases from 0 to 2, such that populations of types 1 and 4 are APC+/+, populations of types 2 and 5 are both APC+/−, and populations of types 3 and 6 are APC−/−.

**Figure1:**
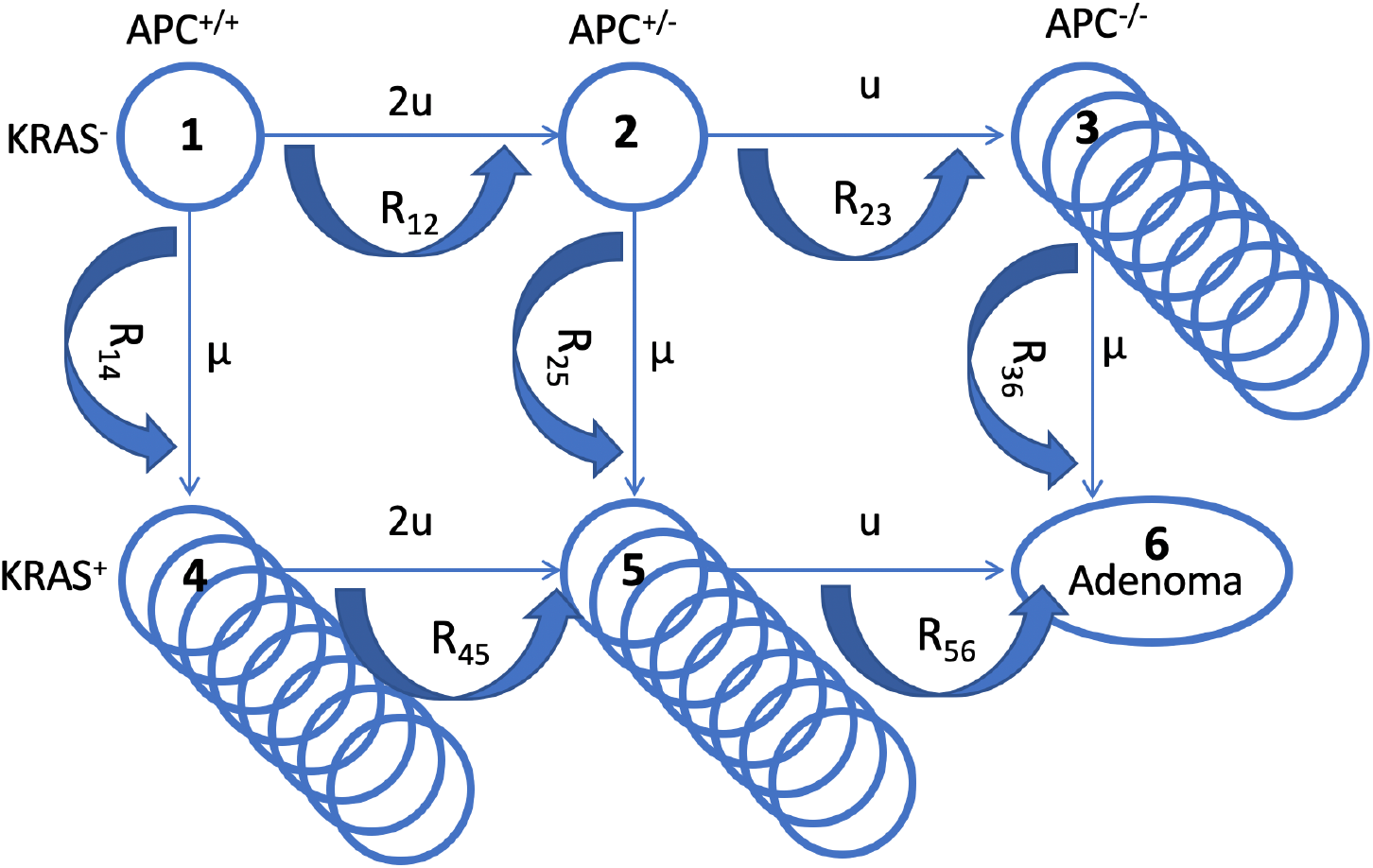
A schematic illustrating the mathematical model. The six cell types are denoted by circles, the mutation rates that give rise to different types are marked by the straight arrows. Crypt conversion rates are indicated by circular arrows and crypt fission by multiple circles.

We model the population dynamics of the colon by using a colonic crypt as a basic unit, which is similar in concept to recently published work [19]. Our model is related to many previous theoretical investigations of the cell population dynamics of crypts [20,21,22,23,24], where stem cells were assumed to acquire random mutations in a constant-population turnover (birth and death) process, and selection happened at the level of individual stem cells. Once it was discovered that there were very few stem cells per crypt [25,26], it became clear that the evolutionary dynamics can be conveniently described at the level of crypts, because crypts are likely to be homogeneous with respect to the driver mutations. The rate at which a crypt changes its mutational status from *i* to *j*, denoted by *R_ij_*, depends on the population size (the number of stem cells), the mutation rate, and the relative fitness of the invading type compared to the resident type [21,22]. The latter can be calculated from the cell displacement data reported in the literature. Types APC+/−, APC−/−, and KRAS+ all have a selective advantage compared to the wild type, which we assume results in an increase of the SC division rate (see the Supplement for details).

Our model keeps track of crypts of different types (denoted as *n_i_* for each type *i*). Modified crypts of types APC−/− and KRAS+ have been reported to undergo crypt fission; in other words, while the total population of a single crypt remains constant (even though it is populated by SCs that are fitter than the wild type SCs), the crypt can undergo a doubling, thus increasing the total number of such modified crypts. The fission rates of different crypt types have been reported in the literature [19,25,27,28] and are denoted by γ_i_; we further denote by δ_i_ the death rate of crypts of type *i*. We model these dynamics by using the following system of ordinary differential equations:

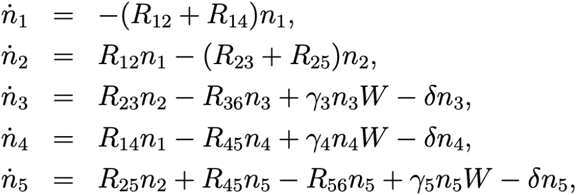

where on the left hand side we have the rate of change for the population of crypts of each type, and W represents competition among modified crypts that undergo crypt fission: W=1-(n_3_+n_4_+n_5_)/K_max_, where K_max_ is the carrying capacity; in reference [19] no crypt competition was included, such that K_max_=∞ and W=1 in their model. The initial conditions for the system above are given by *n*_1_(0) = *N_crypt_*, *n_i_*(0) = 1 ≤ *i* ≤ 5, that is, initially all N_crypt_ crypts are wild type. Parameter values are presented in **Table 2** of the Supplement, and parameter δ=0 unless specified otherwise. In the literature, the APC−/− genotype has been related to the appearance of aberrant crypts (type 3), and an activation of the KRAS oncogene is connected with the growth of polyps (types 4 and 5). In other words, both mutational events are associated with a (pre-)malignant phenotypic change. The combination of both types of mutations (type 6) is thought to correlate with the growth of advanced adenomas. The probability to have produced a crypt of type 6 (i.e. advanced adenoma) is denoted by P(t) and is given by the following equation,

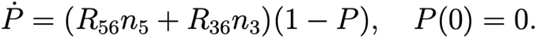

While the model assumptions about the pathways to adenoma formation are clearly defined in our model, it is important to point out that there are uncertainties in those assumptions, and that there is heterogeneity in the types of mutations that can lead to colorectal carcinogenesis. For example, it has been reported that among non-hypermutated colorectal tumors, KRAS was mutated in only about 43% of patient samples [29], indicating the importance of evolutionary pathways that are not captured in our model. Similarly, as in the previous study [19], we assume flexibility regarding the order with which the different mutations can occur. Hence, it is assumed that the initial mutation can occur either in APC or in KRAS. It is, however, controversial whether adenoma formation can indeed be initiated by a mutation in KRAS. Some studies indicate that an initial mutation in KRAS leads to the formation of non-dysplastic polyps, which could represent an evolutionary dead end for neoplasias [30,31]. On the other hand, it has been suggested that an initial KRAS mutation might be able to drive the initiation of colorectal carcinogenesis [32,33], based on mutation frequencies in aberrant crypt foci and adenomas. The assumed flexibility in the evolutionary pathway of the model accommodates these conflicting notions.

### The adenoma incidence data

In order to study the incidence of adenoma, we used the data reported in [34] for the age-ranges 55-59, 60-64, 65-69, 70-74, and 75-79. While this study provides incidence data for nonadvanced adenoma, advanced adenoma, and colorectal cancer (CRC), we focused only on the combined incidence of advanced adenoma and cancer. This assumes that individuals that have developed CRC have most likely already developed an advanced adenoma by the age of testing, and further that nonadvanced adenoma likely refers to fewer mutational steps compared to our type 6, where both the APC gene is fully inactivated and the KRAS gene is mutated. The paper reports data separately for males and females; for our purposes we combined the two values to study the average, since the model is not sufficiently detailed to distinguish between the genders.

## Results

### Fitting the adenoma incidence curve

Until recently, most of the parameters associated with cellular dynamics in colonic crypts were unknown, but presently many of the rates have been estimated with a high degree of confidence [19], which makes it possible to parameterize the model and use it to answer questions about the process of crypt transformation and the dynamics of cancer initiation. Using the published data on the mutation rates, the total number of crypts, the number of SCs per crypt, and the relative fitness of different cell types (see **Table 2 of the Supplement**), we first attempted to fit the model in the absence of crypt competition (W=1), by varying the SC division rate within the physiological range and finding the best fitting value for crypt fission rates. The best fitting parameter combinations always corresponded to zero crypt fission rates. Non-zero crypt fission rates resulted in a much steeper rise in the adenoma incidence compared to that reported in [34]. A similar result was obtained when we used different values for fitness differences (the exhaustive parameter search and a model selection procedure are described in the Supplement). Finally, using the reported crypt fission rates (**Table 2 of the Supplement)** we were not able to find a SC division rate within the biologically applicable range that would give the correct shape of the adenoma incidence curve. The conclusion is that an unlimited exponential expansion of crypts by fission gives an unrealistically steep rise in incidence. This problem did not occur in reference [19] because fitting of the whole incidence of CRC was not attempted, and instead, only the total life-time risk of CRC was compared to the model prediction.

Including crypt competition in the model has resolved this issue. Fixing the carrying capacity parameter K_max_ to a value that is much smaller than the total number of crypts (to ensure that crypt competition significantly restricts the outgrowth of the transformed crypts), we were able to fit the data for a wide range of the SC division rates, with the non-zero best-fitting crypt fission rates that have the correct order of magnitude. Additionally, fixing the crypt fission rates to their reported values, we were able to find very well-fitting incidence curves for a wide range of SC division rates, with the carrying capacity parameter K_max_ ranging between about 500 and about 5000.

For the model that includes crypt competition, it was possible to find nearly equally good fits for a range of biologically plausible parameter values, see **Fig. 2**. The amount of data in the adenoma incidence curve does not allow finding unique values for all the parameters, but instead it allows using many of the parameters fixed to their experimentally obtained values, and just fine-tuning the small number of remaining parameters whose value is unknown (such as K_max_) or only its range is known (such as the SC division rate). When using the parameterized model to study the role of aspirin, instead of selecting the best fitting parameter set, we included best fitting parameter ranges, to see how this variability influences the result.

**Figure 2:**
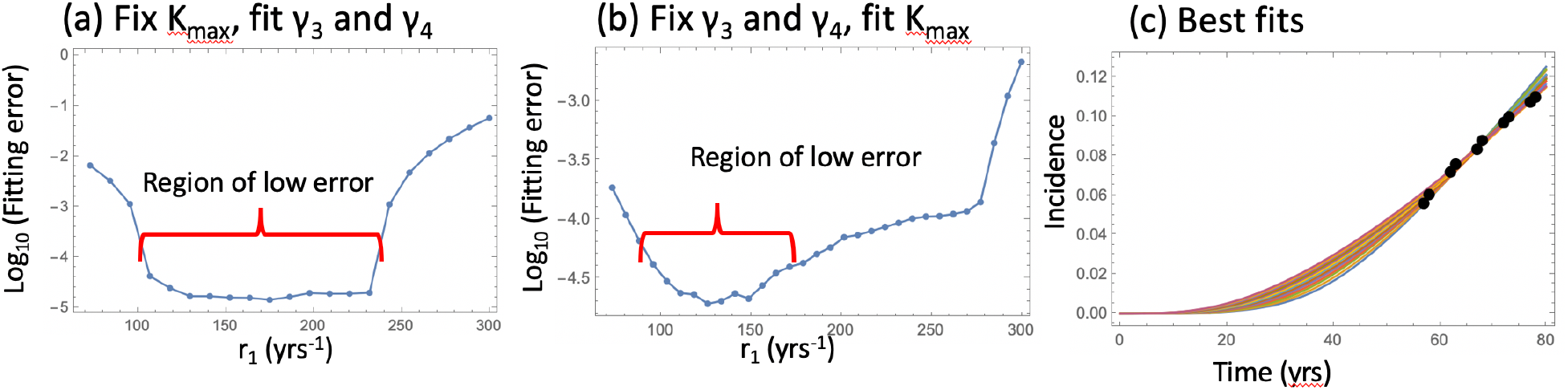
Fitting the nonlinear model to late adenoma incidence data. (a) The fitting error as a function of r_1_, for the best fitting pairs (γ_3_ and γ_4_) with K_max_=1000, and the rest of the parameters are as in table 2 of the Supplement. (b) The fitting error as a function of r_1_, for the best fitting value of K_max_, with the rest of the parameters are as in table 2 of the Supplement. (c) The best fitting curves corresponding to increasing SC division rates, r_1_, are plotted together with the epidemiological data (the values of r_1_ are taken from the Region of low error in panel (b)).

### Pathways to adenoma

Next, we asked what is the most likely pathway that leads to the creation of the type 6 (advanced adenoma). It is possible that crypts of type 6 could be created by a KRAS mutation in a crypt of type 3 (we called this “APC-path”), or by an APC mutation in a crypt of type 5 (“KRAS path”), see panel (c) of figure 3. We found, consistent with [19], that the likelihood of each of these two pathways is determined by the crypt fission rates, and not by mutation rates or crypt conversion rates. Results are presented in **Fig. 3**.

**Figure 3:**
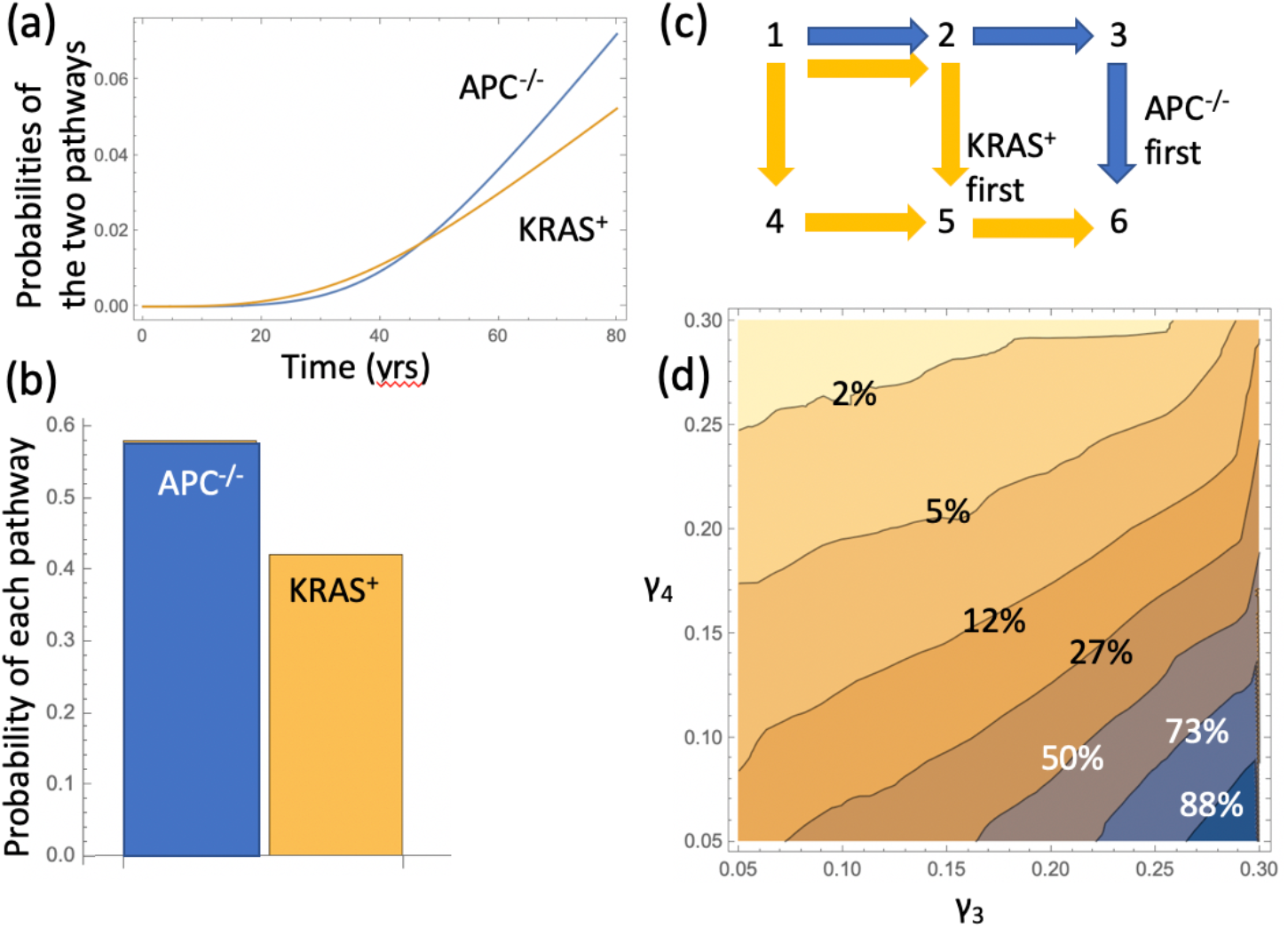
Pathways to adenoma. (a) The probabilities P_APC_ and P_KRAS_ are plotted as functions of time for the best-fitting parameter set of figure 2(b) which corresponds to K_max_=1318 and r_1_=141.1. (b) For the same parameters, the probabilities that adenoma is created by each of the two pathways are shown as bars; the bars represent the quantities P_APC_/(P_APC_+ P_KRAS_) and P_KRAS_/(P_APC_+ P_KRAS_) at t=80 yrs. (c) A schematic representation of the two pathways. (d) The predicted probability of adenoma to be generated through the APC^−/−^ pathway; the quantity plotted is the same as in panel (b) for this pathway. For each pair of rates γ_3_ , γ_4_, the best fitting values of r_1_ and K_max_ were found and the probabilities of the pathways calculated; the rest of the parameters are as in **Table 2 of the Supplement.**

The model allows for the calculation of the probabilities to develop an advanced adenoma through the APC−/− and KRAS pathways, functions P_APC_(t) and P_KRAS_(t). Panel (a) of figure 3 plots these quantities as functions of age (t) for the best fitting parameters of figure 2(b) (K_max_= 1318, r_1_ =141.1). We observe that after the age of about 50, the pathway through the inactivation of the APC gene becomes predominant, with just under 60% of all adenomas at the age of 80 years created by the APC^−/−^ first pathway (panel (b)). This is consistent with the conclusions of reference [19].

We have also investigated the prevalence of the APC−/− path more generally, to see how it depends on the relative values of the fission rates of APC^−/−^ and KRAS^+^ crypts (parameters γ_3_ and γ_4_ respectively). Panel (d) shows a heat plot of the probability of the adenoma (at age 80) to be created through the APC^−/−^ pathway; here the darker colors correspond to a higher likelihood of the APC-path relative to the KRAS-path. We can see that if the fission rate of APC−/− crypts is higher than that of KRAS+ crypts, then the APC-first pathway is more likely (the right bottom corner of the heat plot in panel (d)). Since the crypt fission rates for APC−/− crypts have been reported to be significantly higher than those for KRAS+ crypts, we conclude that the inactivation APC gene is likely to be the first genetic event leading toward advanced adenoma.

### The effect of aspirin

We asked, given that a variety of parameter values could lead to the same incidence curve, can we still say anything about the possible role of aspirin in cancer prevention/delay? To model the effect of aspirin on the relevant kinetic parameters, we used a variety of sources. One type of data was obtained by us in our earlier studies, where the effect of aspirin was quantified by measuring cells’ kinetic parameters with and without aspirin treatment, in vitro and in xenografts [7,8]. In other work, it has also been demonstrated that a related non-steroidal anti-inflammatory drug, sulindac, inhibited the fission of *Apc*-deficient crypts and thus reduced adenoma numbers in mice.

It is, however, unclear which exact cell populations aspirin might affect in vivo. Therefore, we implemented the effect of aspirin in the epidemiological model by testing three different sets of assumptions: (a) the fitness of type 6 cells is reduced; (b) the fitness of type 2-6 cells is reduced; (c) crypt fission rate is reduced.

The first two models (see (a) and (b) below) assume that aspirin reduces the fitness of some cell types. The effect on fitness could be the result of a reduction of the division rate, an increase in the death rate, and/or an increase in the differentiation rate of the cells; for the purposes of our model, it is the combined effect that changes the cells' relative fitness and decreases the probability of crypt conversion. For example, in our previous in vitro study [8], the strongest aspirin dose was associated with a (roughly) two-fold decrease in the division rate of cells and a roughly 1.5-fold increase in the death rate of cells. In our in vivo study [7], parameter changes under different aspirin doses were measured in xenografts. It was found that the strongest dose (100 mg/kg) resulted in a roughly 35% reduction of the cell division rates and a roughly two-fold increase in the death rate. A smaller dose of 15 mg/kg was associated with an approximately 14% reduction in the cells’ division rate and a 30% increase in the death rate.

Translating this information into the fold decrease in SC fitness is not a straightforward task. In particular, while fold-reduction in division rate could be directly implemented, an increase in death rate is less straightforward. This is because of differences between the SC dynamics in healthy or partially transformed colonic crypts modeled here, and cell line dynamics observed in our previous experiments [7,8]. Cell lines undergo a net expansion, which is a result of cell divisions and cell deaths (apoptosis). It was established that the rates of both of these processes were affected by aspirin. On the other hand, the dynamics in colonic crypts is a balance between SC proliferations and SC differentiations, with a possible contribution of SC apoptosis. It is important to note that SC death in the absence of aspirin treatment is not likely a big contributor to the turnover dynamics. Therefore, if the rate of SC apoptosis is increased, say, two-fold in the presence of aspirin, this does not translate to a two-fold reduction in SC fitness. In the extreme scenario of zero SC death in the absence of aspirin, a two-fold increase in this parameter will not lead to a change in SC fitness.

Taking this into account, we view the aspirin-related increase in cell death as a less important factor compared to the decrease in the self-renewal rate. Aspirin-related changes in the self-renewal rate alone can lead to a two-fold decrease in cell fitness (factor of 0.5, associated with the strongest dose of 100 mg/kg); the weakest dose tested (15 mg/kg) leads to a factor of 0.86. Further, if we assume that SC death comprises 10% of all SC “removal” (that is, in the absence of aspirin, the SC apoptosis rate is about 1/10 of SC differentiation rate), then the death rate increase by a factor of 2 (the largest observed) translates into a 10% increase in the removal rate, or an additional factor of 0.9 multiplying the SC fitness. For the purposes of this study we will therefore focus on the range of aspirin fitness reduction factors between 0.45 for the largest aspirin dose, 0.85 for the dose of 15mg/kg, and higher for smaller doses.

### (a) Aspirin reduces the relative fitness of cells of type 6

(that is, the most modified cell type that combines both the APC−/− mutation and the KRAS+ mutation). Assuming that aspirin reduces the relative fitness of type 6, we postulate that the crypt conversion rates R_36_ and R_56_ are modified by a factor less than one. Using the best fitting parameter combination from figure 2(b), we reduced the fitness of type 6 cells by multiplying it by a factor from 1 to 0; the resulting incidence curves are shown in the top plot of panel (a) of figure 4: the stronger the fitness reduction, the lower the incidence. The bottom graph shows the relative incidence at age 70, which is defined as the incidence under aspirin treatment divided by the incidence in the absence of aspirin treatment. This quantity for advanced adenma has been reported to vary between about 0.5 (the dashed horizontal line) and 1, depending on the aspirin dose [35]. The relative incidence is plotted as a function of the aspirin-induced factor that reduces the fitness of the type 6 cells; as expected, the relative incidence is a monotonic function of this factor. Instead of plotting a single curve corresponding to the best fitting parameters of figure 2, we plotted several curves that correspond to the region of low error in that figure. We observe that these fits (which correspond to different values of SC division rate) result in very similar relative incidence plots.

We can compare the obtained relative adenoma risk with the data reported in the literature, see e.g. [35,36,37,38]. In particular, the dose-dependence of colorectal adenoma was studied [35], and it was shown that the relative risk for adenoma was 0.80 for women who used 0.5 – 1.5 standard tablets per week, 0.74 for those who used 2 – 5 tablets per week, 0.72 for those who used 6 – 14 tablets per week, and 0.49 for those who used more than 14 tablets per week. Comparing this with the relative adenoma risk plot in figure 4(a), we can see that the model predictions are very consistent with the observed bounds: for the strongest aspirin dose, which translates in the fitness reducing factor of 0.45, the relative risk is predicted to be just under 0.5, and for the dose of 15 mg/kg (factor of 0.85), the relative risk is about 0.85.

### (b) Aspirin reduces the relative fitness of cells of types 2—6

(that is, all the modified types). The results are presented in panel (b) of figure 4. As in the previous case (panel (a)), different values of SC division rate lead to very similar relative incidence curves. Since not only rates R_36_ and R_56_ are now lowered, but also rates R_12_ and R_14_, the incidence of adenoma decreases more, which results in a lower value for the relative incidence curve (the bottom graph). For example, the highest aspirin dose (which corresponds to the factor of 0.45 on the horizontal axis) is predicted to lead to a more than 5-fold reduction in the adenoma incidence, which is a much stronger reduction than two-fold, as observed in [35]. Therefore, we conclude that in the framework of this model aspirin is not likely to strongly reduce the fitness of partially-transformed cells, or that its influence on such cells is significantly weaker compared to the fitness reduction of the type-6 cells (advanced adenoma cells).

### (c) Aspirin reduces the fission rate of the crypts

by lowering the values of parameters γ_3_ and γ_4_, (see panel (c) of figure 4). As before, the incidence of adenoma is predicted to be reduced under aspirin, but compared to the previous simulations, the exact shape of the relative incidence function is more sensitive to the base SC division rate. In other words, even though the parameter combinations from the Region of low error in figure 2 all produce very similar fits to the incidence curve, the incidence reduction due to aspirin is sensitive to these parameters. As for the extent of the aspirin-induced reduction in the adenoma risk, it appears to be consistent with the reported risk reduction at least for a subset of the parameter combinations. Therefore, this mechanism cannot be rejected based on the predicted late adenoma incidence reduction.

**Figure 4.**
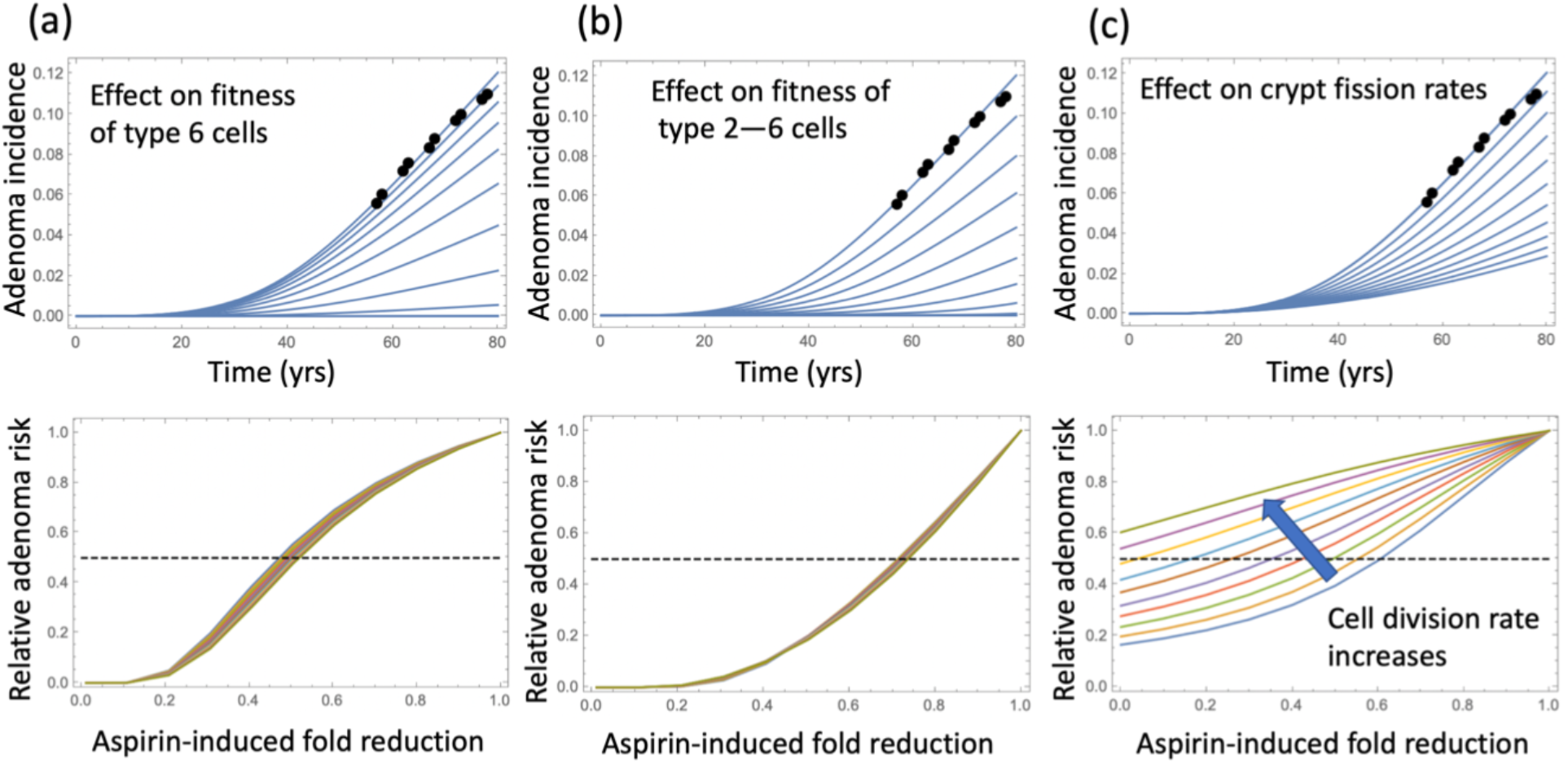
The effect of aspirin on the adenoma incidence curve. Top row: adenoma incidence curves are plotted for the best fitting parameter combination of figure 2(b); the different curves correspond to the different degrees of aspirin influence. Bottom row: the relative incidence at age 70 plotted as a function of the aspirin-induced fold reduction of the appropriate quantity, which is (a) fitness of type 6 cells, (b) fitness of type 2,3,4,5,6 cells and (c) crypt fission rate. The relative fitness is plotted for all the parameter combinations in the region of low error in figure 2(b). The rest of the parameters are as in figure 2(b).

## Discussion

We used mathematical modeling approaches to test the hypothesis that the changes in tumor cell kinetics observed during aspirin treatment in vitro and in vivo can translate into a protective effect on a population level that is consistent with epidemiological observations for late adenoma. This was done by first constructing a mathematical model of in vitro carcinogenesis describing evolutionary events leading up to the late adenoma stage. This model was then used to calculate expected population incidence as a function of age. Many of the model parameters have recently been estimated experimentally, which provides a solid basis for this modeling effort. Remaining parameters were estimated by fitting the incidence prediction to epidemiological data on late adenoma detection. A linear model that did not include inter-crypt competition was rejected because its best fits corresponded to zero crypt fission rates, and the more (statistically) powerful model was adopted instead, where individual mutated crypts experienced both fission and nonlinear competition dynamics. This parameterized model was used as a basis to explore how changes in the kinetics / fitness of cells, brought about by aspirin, can modify the predicted incidence of late adenomas.

Both our in vitro and in vivo work indicated that aspirin reduces the rate of colorectal tumor cell division and increased the rate of tumor cell death in a dose-dependent way, by up to two-fold for the largest aspirin dose used in these experiments. Our modeling in the current study has demonstrated that changes in the cellular fitness of a magnitude that lies within our experimentally observed range can lead to significant reductions in late adenoma incidence. Epidemiological data suggest that adenoma incidence is reduced by aspirin in a dose-dependent manner, with the incidence cut in half for the highest doses explored [35]. A reduction of this magnitude is predicted in our model if aspirin is assumed to reduce the cellular fitness generally less than two-fold, which is within our experimentally observed range. Therefore, we can conclude that the aspirin-induced changes in cellular fitness that we observed experimentally can in principle explain a significant portion of the protective effect observed on the population level.

This does of course not preclude alternative mechanisms that can further contribute to the protective effect. It is very likely that a reduction in the level of inflammation within the microenvironment of the cells can reduce the incidence of colorectal cancer, because inflammation has been identified as a driver of this disease [6]. Furthermore, the CAPP2 study [5] showed that the protective effect of aspirin was only observed after a follow-up time of more than 55 months, indicating that further, yet to be determined, complexities are at work that lead to this delay in outcome. Our analysis, however, points out that a direct effect of aspirin on the tumor cells can be one important determinant of protection.

As with most mathematical modeling studies, there are uncertainties in assumptions that need to be kept in mind. Our experiments [7,8] were performed with tumor cell lines, both in vitro and in mouse xenografts. Cellular processes in the human colon, however, are most likely driven by stem cell dynamics, and differences in response to aspirin may exist. Data from mice, however, indicate that stem cells react to aspirin in a similar way as documented in our experimental studies [39], indicating that these effects carry over. Another point of uncertainty concerns the identity of the cell populations that are affected by aspirin. To address this, we made several assumptions, and results remained robust. Thus, we assumed that aspirin influences only the most advanced adenoma cell population, characterized by APC−/− and KRAS+ mutations. Results remained fairly similar in an alternative model, where all cells that have either acquired an APC or a KRAS mutation are impacted by aspirin (although in this case the effect of aspirin is stronger, which is not surprising given that a larger cell population loses fitness). There is evidence that aspirin might influence not only the cell dynamics themselves, but also the crypt fission dynamics, reducing the rate at which crypts divide. Incorporation of this effect into our model does not lead to qualitative changes in our conclusions.

An important component of all of this work is the underlying mathematical model of in vivo adenoma formation. The assumptions about the genetic events that occur during adenoma formation are consistent with our current understanding of adenoma evolution [17], and a similar model that also includes evolutionary events beyond adenomas has recently been published [19]. An important difference between our and the previous model concerns assumptions about crypt fission dynamics. The previous study [19] assumed that crypt fission can occur without density-dependent effects. Using experimentally available parameter estimates, this model could account for the life-time risk of colorectal cancer. When applying a similar model to late adenoma age incidence data, however, we could not obtain a good fit for the age-incidence curve, and the best fit was in fact obtained in the absence of any crypt fission. With unlimited crypt fission, the predicted adenoma incidence rose too sharply with age compared to epidemiological data. When introducing density-dependence into the crypt fission process, however, late adenoma age incidence data could be readily fit, and so we used this model assumption to go forward. Indeed, it is likely that density-dependent effects play a role in crypt fission, because this process is probably influenced by signaling factors that become limiting as the number of crypts increases. It would be important to verify this assumption experimentally in future work.

Crypt fissions is a process that is well documented with data [26], and this is the reason that we incorporated it into our model. The question can, however, be asked whether crypt fission is the primary driver of transformed cell expansion, or whether a more general proliferation of cells, with consequent development of altered crypts or glands, drives disease development. It turns out that similar results are obtained when a model is considered assuming that density-dependent proliferation of cells beyond the crypt size (rather than proliferation of crypts) drives adenoma development. This model is presented in the Supplementary materials, and results in similar conclusions when used to quantity the effect of aspirin.

In conclusion, this modeling analysis suggests that a direct impact of aspirin on the kinetics and fitness of mutated cells can significantly reduce the incidence of colorectal adenomas, with a magnitude that is consistent with epidemiological data. This highlights the importance of investigating this effect of aspirin experimentally in more detail, especially under experimental conditions that approximate cell dynamics in the human colorectal tissue with greater accuracy.

## Acknowledgements

Support of the following grants is gratefully acknowledged: NIH 1 U01 CA187956-01 (AG, RB, NK, DW); NSF-Simons Center for Multiscale Cell Fate Research (NK, YW); NIH/NCI U54-CA217378 (NK, DW, YW).

