## Supplementary Materials for "The protective effect of aspirin in colorectal carcinogenesis: a multiscale computational study from mutant evolution to age incidence curves"

### 1 A crypt-based model of adenoma initiation

#### 1.1 Mathematical formulation

Let us enumerate the types in the way presented in table 1.

| Mutations in APC | Mutations in KRAS | Type number |
|---|---|---|
| 0 | 0 | 1 |
| 1 | 0 | 2 |
| 2 | 0 | 3 |
| 0 | 1 | 4 |
| 1 | 1 | 5 |
| 2 | 1 | 6 |

Table 1: Enumeration of the different genotypes.

Then we can denote by $n_i$ with $1 \leq i \leq 6$ the number of crypts of type $i$. Suppose $R_{ij}$ is the conversion rate from crypt type $i$ to type $j$, and $\gamma_i$ the growth rate (by crypt fission) of crypts of type $i$. We have the following equations:

$$\begin{aligned} \dot{n}_1 &= -(R_{12}+R_{14})n_1, & (1)\\ \dot{n}_2 &= R_{12}n_1-(R_{23}+R_{25})n_2+\gamma_2 n_2, & (2)\\ \dot{n}_3 &= R_{23}n_2-R_{36}n_3+\gamma_3 n_3, & (3)\\ \dot{n}_4 &= R_{14}n_1-R_{45}n_4+\gamma_4 n_4, & (4)\\ \dot{n}_5 &= R_{25}n_2+R_{45}n_5-R_{56}n_5+\gamma_5 n_5, & (5)\end{aligned}$$

with the initial conditions

$$n_1(0)=N_{crypt}, \quad n_i(0)=0, \quad 1\le i\le 5. \tag{6}$$

In the ODEs above, we have ignored the process of stochastic tunneling such that the crypts can only convert one step at a time. It is further possible to ignore the negative (outgoing) rates, which simplifies this linear system to the following:

$$\begin{aligned} \dot{n}_1 &= 0, & (7)\\ \dot{n}_1 &= R_{12}n_1+\gamma_2 n_2, & (8)\\ \dot{n}_3 &= R_{23}n_2+\gamma_3 n_3, & (9)\\ \dot{n}_4 &= R_{14}n_1+\gamma_4 n_4, & (10)\\ \dot{n}_5 &= R_{25}n_2+R_{45}n_5+\gamma_5 n_5. & (11)\end{aligned}$$

The probability $P(t)$ that by time $t$ at least one crypt of type 6 has been created is given by the solution of the equation

$$\dot{P}=(R_{56}n_5+R_{36}n_3)(1-P), \quad P(0)=0. \tag{12}$$

The solution can be obtained exactly and is given by

$$P=1-exp\left\{-N_{crypt}(S_{45}R_{1\to4\to5\to6}+S_{25}R_{1\to2\to5\to6}+S_{23}R_{1\to2\to3\to6})\right\}, \tag{13}$$

where the quantities $S_{ij}$ correspond to the paths $1\to i\to j\to 6$,

$$R_{1\to i\to j\to 6}=R_{1i}R_{ij}R_{j6}, \tag{14}$$

see figure 1. They can be written down by using the following function:

$$g(x)=\frac{e^{xt}-1-xt}{x^2}>0. \tag{15}$$

We have

$$g(0)=\frac{t^2}{2}, \quad g'(x)\equiv\frac{\partial g}{\partial x}=\frac{2+tx+e^{tx}(tx-2)}{x^3}>0,$$

that is, this function increases monotonically in $x$. We have

$$S_{ij}=\frac{g(\gamma_i)-g(\gamma_j)}{\gamma_i-\gamma_j}.$$

Note that expressions $S_{ij}$ have a singularity if any of the quantities $\gamma_i$, $\gamma_j$ is zero and/or if $\gamma_i = \gamma_j$. For example, if both growth rates are zero ($\gamma_i = \gamma_j = 0$), we have

$$S_{ij} = \frac{t^3}{6}.$$

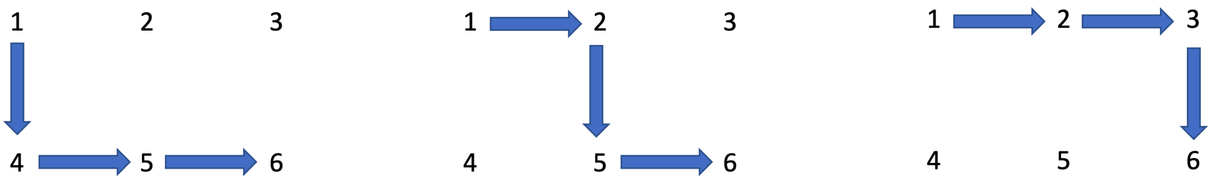

Figure 1: Three pathways to adenoma.

In our context it is reasonable to assume that $\gamma_2 = 0$, that is, APC$^{+/-}$ crypts do not divide, and that $\gamma_4 = \gamma_5$, that is, APC$^{-/-}$ and APC$^{+/-}$ crypts with an additional KRAS mutation divide at the same rate. In the $\gamma_i = \gamma_j$ case, taking the limit as $\gamma_j \to \gamma_i$, we obtain

$$S_{45} = g'(\gamma_4), \quad S_{25} = \frac{g(\gamma_4) - t^2/2}{\gamma_4}, \quad S_{23} = \frac{g(\gamma_3) - t^2/2}{\gamma_3}. \tag{16}$$

It is convenient to view $S_{25}$ as a function of $\gamma_4$, $S_{25} = F(\gamma_4)$, where

$$F(x) = \frac{1}{x^3}\left(e^{xt} - \left(1 + xt + \frac{(xt)^2}{2}\right)\right) = \frac{1}{x^3}\sum_{i=3}^{\infty}\frac{(xt)^i}{i!},$$

then $S_{23} = F(\gamma_3)$, and $F(x)$ is an increasing function of $x$. That is, if $\gamma_4 > \gamma_3$ then $S_{25} > S_{23}$, and if $\gamma_4 < \gamma_3$ then $S_{25} < S_{23}$. We further note that $S_{45} = \tilde{F}(\gamma_4)$ with

$$\tilde{F}(x) = \frac{1}{x^3}\left(e^{xt}(tx - 2) + tx + 2\right) = \frac{1}{x^3}\sum_{i=3}^{\infty}\frac{(xt)^i(i-2)}{i!} > F(x).$$

To summarize, we can see that $S_{45} > S_{25}$, and $S_{23}$ is greater (slammer) than $S_{25}$ if $\gamma_3$ is greater (smaller) than $\gamma_4$.

The conversion rates are given by

$$R_{ij} = r_i K u_{i\to j} \rho_{ij}, \tag{17}$$

where $r_i$ is the division of type $i$, $u_{i\to j}$ is the mutation rate from type $i$ to type $j$, and $\rho_{ij}$ is the probability that one cell of type $j$ becomes fixated in a compartment of size $K$ with the host type $i$. To calculate this probability, let us denote by $d_i$ the death rate of type $i$. Then we have

$$\rho_{ij} = \frac{1 - \frac{r_i d_j}{r_j d_i}}{1 - \left(\frac{r_i d_j}{r_j d_i}\right)^K}, \tag{18}$$

where $\frac{r_i d_j}{r_j d_i}$ is the inverse of the relative fitness of type $j$ with respect to type $i$.

#### 1.2 Fitting the linear model to late adenoma incidence data

The model to fit to the adenoma incidence curve is given by equations (13,14,15,16,17,32). We will make the following assumptions (see also table 2):

- Wild-type crypts and those with only a single copy of APC gene mutated do not proliferate ($\gamma_1 = \gamma_2 = 0$).
- Crypts with a KRAS mutation and $APC^{+/+}$ and $APC^{+/-}$ phenotypes proliferate at the same rate, $\gamma_4 = \gamma_5$.
- The fitness of cells with $APC^{+/-}$, $APC^{-/-}$, and $KRAS^+$ phenotypes relative to the wild type cells was determined using the cell replacement data from [6]. In particular, we assumed that for any phenotype,

$$F_j \equiv \frac{r_j d_1}{d_j r_1} = \frac{Pr(j)}{1 - Pr(j)}, \quad j \in 2, 3, 4$$

  where $Pr(j)$ is the probability of replacement of the wild type by type $j$ found in [6]. Using this formula, we obtain the values given in table 2. Then the fitness of other types is multiplicative, $F_5 = F_4 F_2$.
- The death rates are assumed the same among the types, such that the division rates of cells satisfy $r_2 = F_2 r_1, r_3 = F_3 r_1$, etc.
- The other parameters are specified in table 2.

| Parameter | Notation | Value/Range |
|---|---|---|
| Number of crypts | $N_{crypt}$ | $10^7$ |
| Number of SCs per crypt | $K$ | 7 |
| Rate of inactivation of APC (per cell division) | $u$ | $10^{-7}$ |
| Rate of inactivation of APC (per cell division) | $\mu$ | $10^{-9}$ |
| Division rate of WT SCs (per year) | $r_1$ | $(18, 365)$ |
| Relative fitness of $APC^{+/-}$ cells | $F_2 = F_{APC+/-}$ | 1.6 |
| Relative fitness of $APC^{-/-}$ cells | $F_3 = F_{APC-/-}$ | 3.76 |
| Relative fitness of $KRAS^+$ cells | $F_4 = F_{KRAS}$ | 3.54 |
| Division rate of $APC^{-/-}$ crypts (per year) | $\gamma_3$ | 0.2 |
| Division rate of $KRAS^+$ crypts (per year) | $\gamma_4$ | 0.07 |

Table 2: Parameters, notations, and their values.

In order to fit the data, we fix parameters $N_{crypt}, K, u, \mu$, and vary the remaining parameters. This is done in stages. We first fix the fitness parameters $F_2, F_3, F_4$ to their values in table (2) and vary the remaining parameters $r_1, \gamma_3, \gamma_4$ to find the global minimum of the error between the model and the data, see figure 2(a-c). Then we take other select values for the relative fitness parameters to show that the results remain qualitatively similar (not shown). Next, we describe the results of fitting and the patterns that were observed.

To find the global minimum of the error, we varied parameter $r_1$ (the division rate of the wild type SCs) between once a day and once every 20 days (which corresponds to the division rate of 365 yrs$^{-1}$ and 18 yrs$^{-1}$). For each value of $r_1$, the error was minimized in the 2-dimensional parameter space $(\gamma_3, \gamma_4)$, and a unique minimum was always found. The best fits corresponding to a subset of these division rates are plotted in figure 2(a), with the best fitting values of $\gamma_3$ and $\gamma_4$ shown in panel (b) for each $r_1$. We can see that as $r_1$ increases, that is, SCs are assumed t divide faster, the best fitting crypt division rates decrease (that is, crypt fission proceeds at a slower rate). For $r_1$ greater than about 250 yrs$^{-1}$ (that is, divisions once every day and a half or faster), the best fitting crypt fission rates are zero. This parameter combination (division rate of about every 1.5 days and zero crypt fission) corresponds to the minimal error of fitting (panel (c)). This is evident from the shape of the best fitting incidence curves, $P(t)$, corresponding to different $r_1$ values (panel (a)). For low rates of S division, the fission rates are relatively high, resulting in an curve $P(t)$ that has a steep rise, yielding a large fitting error and a qualitatively unrealistic shape of the incidence curve if compared with the data. This is the consequence of a pronounced exponential increase in the number of crypts, which sharply accelerates the generation of type 6 crypts. As $r_1$ increases and crypt fission rates decrease, the incidence curves become less steep, until the best fitting crypt fission rate reaches zero. At this stage, the best fitting incidence curve is achieved, because further increase in $r_1$ results in an increase in the incidence that happens too early.

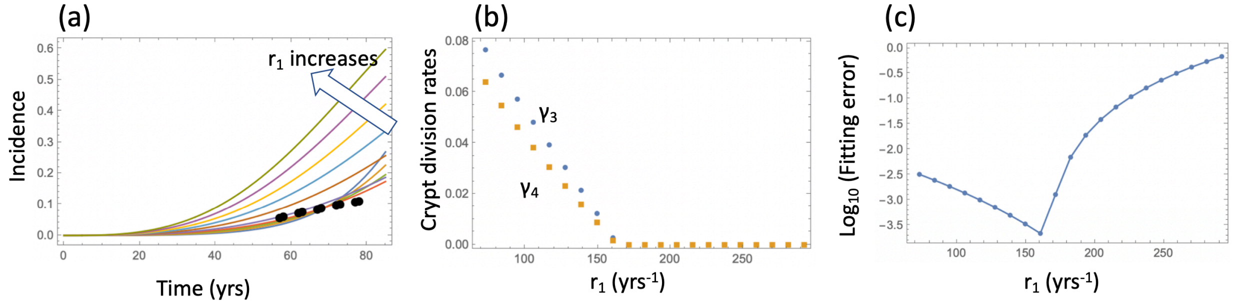

Figure 2: Model fitting to the incidence data. (a) The best fitting curves corresponding to increasing SC division rates, $r_1$, are plotted together with the epidemiological data. (b) The best fitting parameters $\gamma_3$ and $\gamma_4$ are shown for each value of $r_1$. (c) The fitting error as a function of $r_1$. The relative fitness values are fixed to $F_{APC^{+/-}} = 1.6, F_{APC^{-/-}} = 3.76, F_{KRAS} = 3.54$. The rest of the parameters are as in table 2.

So far, in order to fit the model to the adenoma data, we fixed several of the parameters to their measured values and varied the three remaining parameters $(r_1, \gamma_3, \gamma_4)$ within realistic ranges to investigated the error landscape and find global minima. To obtain a more comprehensive picture of the model behavior, we have implemented a procedure where five parameters were varied: the division rate of stem cells, $r_1$, two cellular fitness parameters, $R_{APC^{+/-}}$ and $R_{KRAS}$ (with $F_{APC^{+/+}} = 2F_{APC^{+/-}}$); and two crypt fission parameters, $\gamma_3$ and $\gamma_4$. The rest of the parameters were set to their values in Table 2.

The fitness parameters were varied between 1.1 and 3.5 to match the measured range. For each pair $(R_{APC^{+/-}}, R_{KRAS})$, a fitting procedure identical to that of figure 2 was performed, see figure 3. Each graph corresponds to a unique pair $(R_{APC^{+/-}}, R_{KRAS})$, and the values of these coefficients are indicated on each panel. The horizontal axes of each panel is the stem cell division rate, $r_1$. The green curves show the $\log_{10}$(fitting error), and the black (gray) dots show the best fitting values of $\gamma_3$ ($\gamma_4$), multiplied by 10 for convenience of presentation. If the best fitting crypt fission parameter was negative, then the fitting error shows corresponded to $\gamma_3 = \gamma_4 = 0$. We observe that in all the cases, the best fitting parameter combination is reached when the crypt fission rates become zero, reproducing the result of figure 2, but for a wide parameter range. To observe this more clearly, we also presented the error landscape for the best fitting parameter $r_1$, as a function of $\gamma_3$ and $\gamma_4$ (the horizontal and vertical axes in each panel, respectively). The quantity $\log_{10}$(fitting error) is shown as a heat map, with darker colors corresponding to lower error values. We can see that the lowest error corresponds to the corner $\gamma_3 = \gamma_4 = 0$.

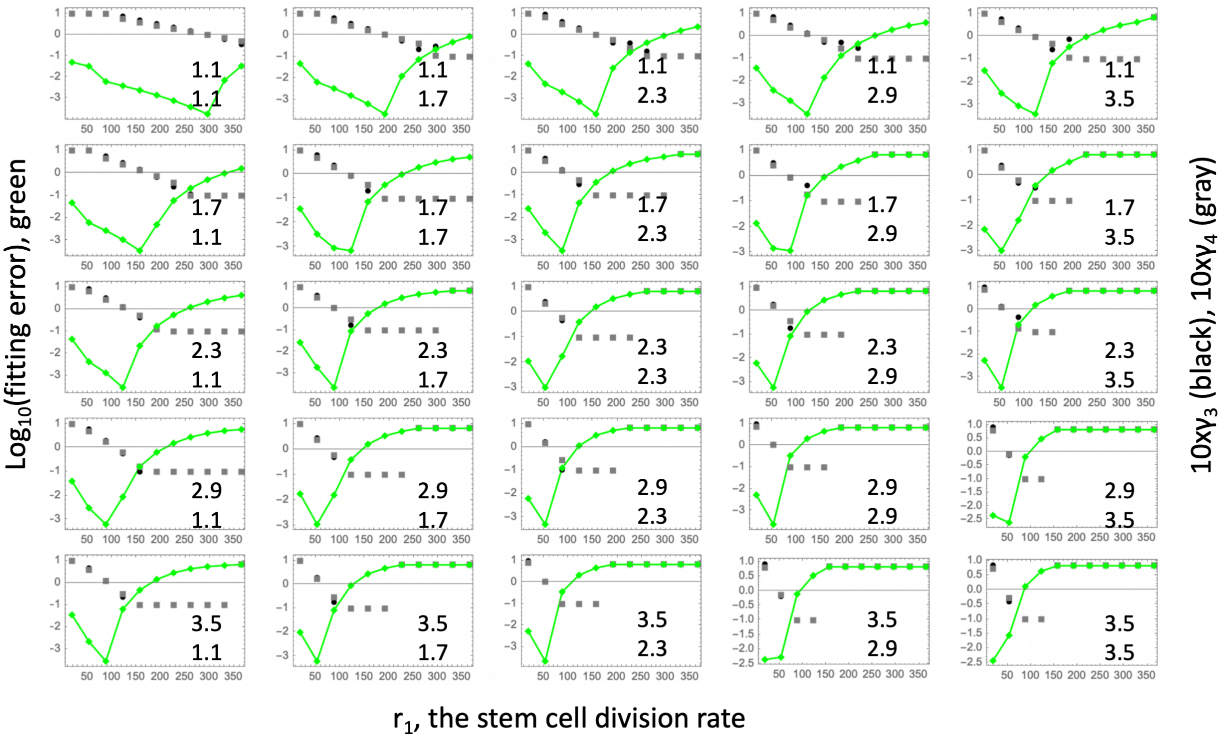

Figure 3: Fitting the linear model. Each panel shows fitting results for a particular parameter combination, $(R_{APC^{+/-}}, R_{KRAS})$; the values of these two fitness parameters are indicated, and $F_{APC^{+/+}} = 2F_{APC^{+/-}}$. The green lines show the $\log_{10}$(fitting error) as a function of $r_1$. The fitted parameters $\gamma_3$ and $\gamma_4$ (multipied by 10) are shown as black (gray) lines. The rest of the parameters are as in table 2.

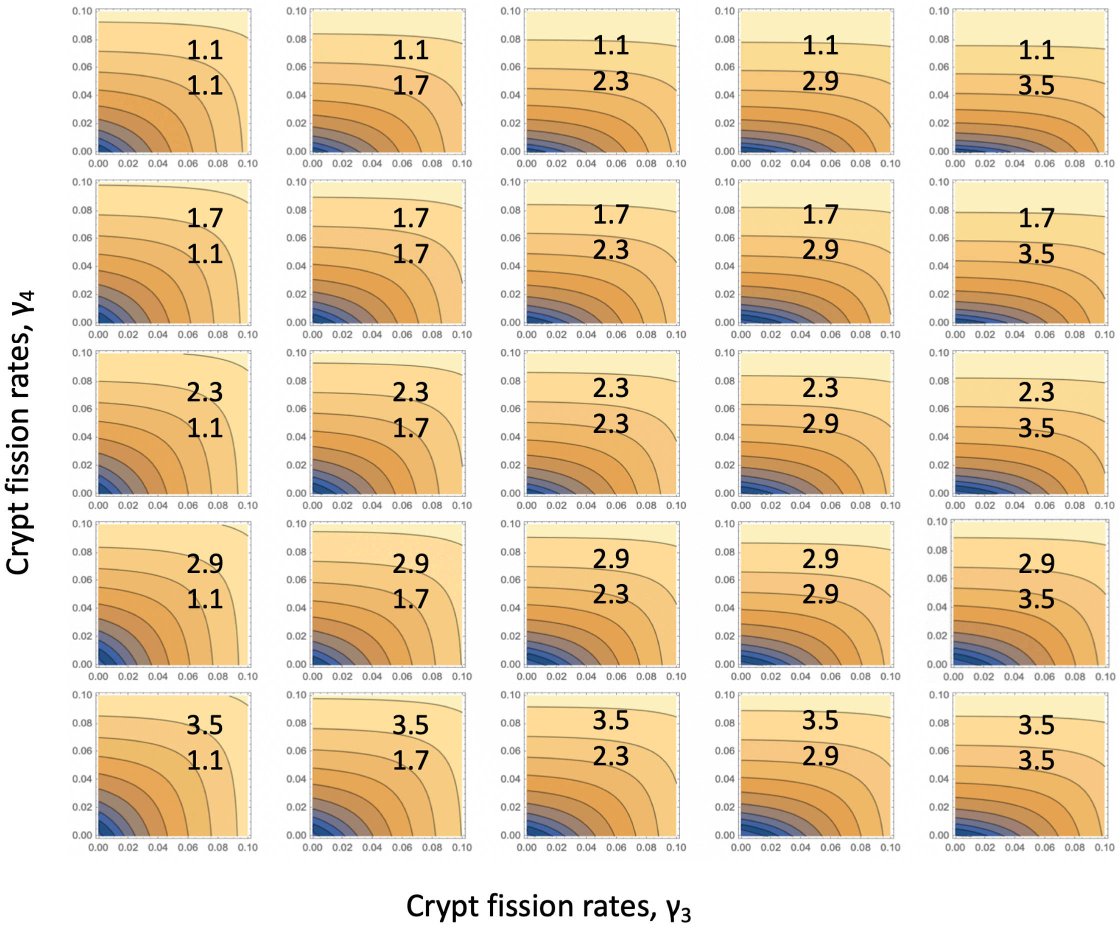

Figure 4: Fitting the linear system: for each parameter combination, $(R_{APC^{+/-}}, R_{KRAS})$, of figure 3, the error landscape is shown that corresponds to the best fitting $r_1$ (see the minima of the green lines in figure 3). The contour plot represents $\log_{10}$(fitting error) as a function of $\gamma_3$ and $\gamma_4$ (darker colors correspond to lower values). The rest of the parameters are as in table 2.

#### 1.3 Nonlinear model: competition among crypts

Let us assume that the types of crypts that undergo crypt fission (types 3,4, and 5) are in direct competition with each other, modeled as logistic, as opposed to straight exponential, growth, under a carrying capacity, $K_{max}$. Let us denote by $W$ the term

$$W = 1 - \frac{n_3 + n_4 + n_5}{K_{max}}.$$

Then the nonlinear system with crypt competition becomes

$$\dot{n}_1 = -(R_{12} + R_{14})n_1, \quad (19)$$

$$\dot{n}_2 = R_{12}n_1 - (R_{23} + R_{25})n_2, \quad (20)$$

$$\dot{n}_3 = R_{23}n_2 - R_{36}n_3 + \gamma_3 n_3 W - \delta n_3, \quad (21)$$

$$\dot{n}_4 = R_{14}n_1 - R_{45}n_4 + \gamma_4 n_4 W - \delta n_4, \quad (22)$$
$$\dot{n}_5 = R_{25}n_2 + R_{45}n_5 - R_{56}n_5 + \gamma_5 n_5 W - \delta n_5, \quad (23)$$

with initial conditions (6) and the probability to create a crypt of type 6 given by system (12). While an analytical solution is no longer available, a procedure similar to that performed for the linear system can be performed numerically. Results can be seen in figure 5. As before, all the parameters were fixed to their values in table 2, except the cell division rate, $r_1$, and the crypt fission rates, $\gamma_3$ and $\gamma_4$. Additionally, we assumed a crypt carrying capacity $K_{max} = 10^3$ and crypt death rate $\delta = 0.05\text{yrs}^{-1}$. The best fitting values of $\gamma_3$ and $\gamma_4$ were found for each value of $r_1$, which was varied in the realistic range. This procedure yielded a range of low-error fits (panel (c)) that correspond to nonzero values of crypt fission rates in the realistic range (panel (b)), with the corresponding incidence curves given in the inset of panel (a).

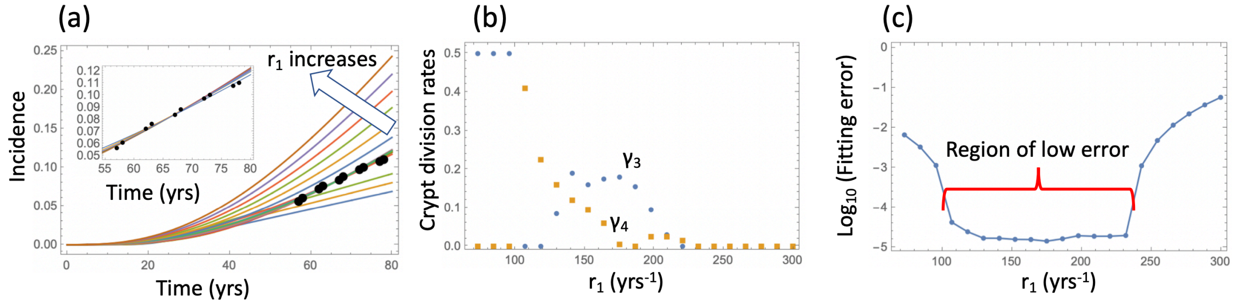

Figure 5: Model fitting to the incidence data, in the presence of crypt competition. (a) The best fitting curves corresponding to increasing SC division rates, $r_1$, are plotted together with the epidemiological data; inset: the best fitting curves corresponding to the values of $r_1$ from the Region of low error, see panel (c). (b) The best fitting parameters $\gamma_3$ and $\gamma_4$ are shown for each value of $r_1$. (c) The fitting error as a function of $r_1$. The relative fitness values are fixed to $F_{APC^{+/-}} = 1.6, F_{APC^{-/-}} = 3.76, F_{KRAS} = 3.54, K_{max} = 1000$. The rest of the parameters are as in table 2.

Since this procedure yielded a wide range of similarly good fits under a fixed value for the crypt carrying capacity, we next performed a fitting procedure where the crypt fission rates $\gamma_3$ and $\gamma_4$ were fixed to those in table 2, and the best carrying capacity, $K_{max}$, was determined for each division rate, $r_1$, by fitting to the data. Results are presented in figure 6. We can see that for a range of values of the cell division rates ,$r_1$, a low-error fit was found, with the carrying capacity values ranging between about $5 \times 10^2$ and $5 \times 10^3$. The best fitting values are $K_{max} = 1318, r_1 = 141.1$ (divisions approximately every 2.5 days). Changing the crypt death rate to $\delta = 0$ yields very similar results, see figure 7.

These results suggest that biologically, the nonlinear model is a more appropriate choice because it produces the best fit for values of crypt fission rates that are within the experimentally observed range, while the linear model requires zero crypt fission rates. From the statistical point of view, the nonlinear model is a significantly more powerful model e.g. by applying the Akaike Infromation Criterion (AIC).

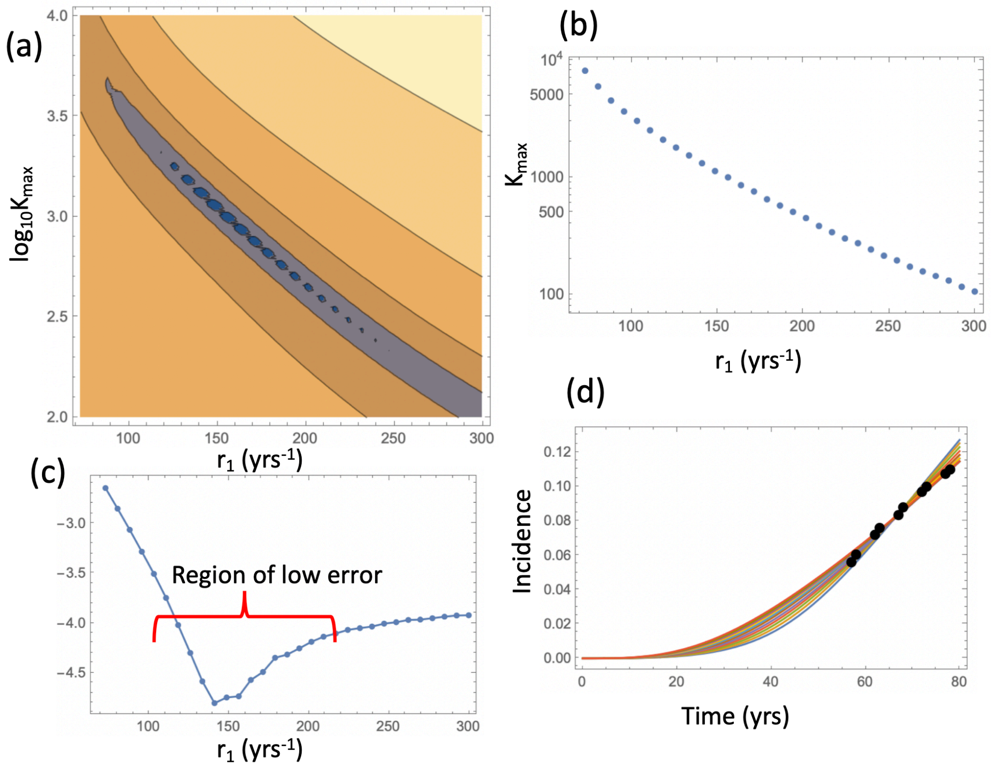

Figure 6: In the presence of crypt competition: the best fitting carrying capacity. (a) The heatplot of the fitting error, where all parameters are fixed except $K_{max}$ and $r_1$. Dark colors correspond to lower values of the error. (b) The best fitting carrying capacity $K_{max}$ is shown for each value of $r_1$. (c) The fitting error as a function of $r_1$. (d) The best fitting curves corresponding to increasing SC division rates, $r_1$, are plotted together with the epidemiological data (the value of $r_1$ are taken from the Region of low error, see panel (c)). The parameter values are fixed to $F_{APC^{+/-}} = 1.6, F_{APC^{-/-}} = 3.76, F_{KRAS} = 3.54, \gamma_3 = 0.2, \gamma_4 = 0.07$, $\delta = 0.05$, and the rest as in table 2.

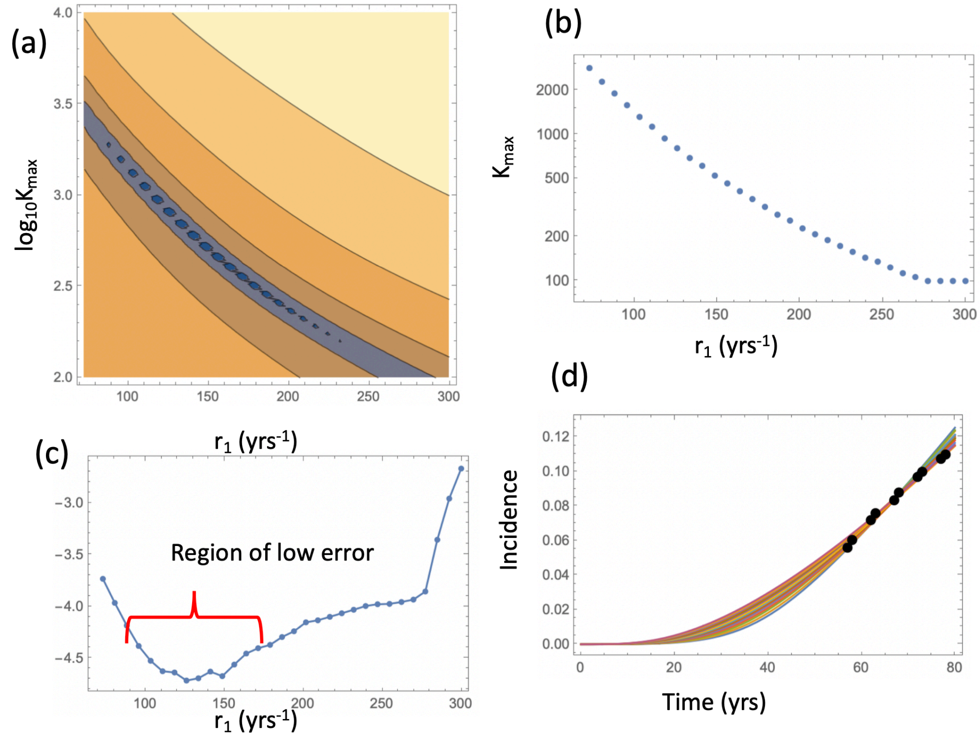

Figure 7: Same as figure 6 but with $\delta = 0$.

In order to determine the likelihood of type $y_6$ produced from cells of type $y_3$ or cells of type $y_5$, we looked at the probabilities of the two pathways, $P_{APC}$ and $P_{KRAS}$, which stand for the probability to create adenoma by first inactivating the APC gene and then adding a gain-of-function mutation in KRAS gene, or by first activating KRAS and then inactivating the APC gene, see panel (c) of figure 3 of the main text. The two probabilities satisfy the following equations,

$$\dot{P}_{APC} = R_{36}n_3(1 - P_{APC}), \quad P_{APC}(0) = 0, \tag{24}$$

$$\dot{P}_{KRAS} = R_{56}n_5(1 - P_{KRAS}), \quad P_{KRAS}(0) = 0. \tag{25}$$

The functions $P_{APC}$ and $P_{KRAS}$ for the best fitting pair of figure 6 ($K_{max} = 1318, r_1 = 141.1$) are presented in panel (a) of figure 3 of the main text. We can see that after the age of about 50, the pathway through the inactivation of the APC gene becomes predominant, with just under 60% of all adenomas at the age of 80 years created by the APC$^{-/-}$-first pathway (panel (b)). In general, the prevalence of each pathway depends on the relative value of the $\gamma_3$ and $\gamma_4$, the fission rates of APC$^{-/-}$ and KRAS$^+$ crypts respectively. Panel (d) shows a heat plot of the probability of the adenoma (at age 80) to be created through the APC$^{-/-}$ pathway.

### 2 A model with cellular expansion

Here we present an alternative modeling concept for the dynamics of cell divisions, mutation and selection (through competition) in a colonic crypt. As before we assume that mutations gave cells fitness advantage, which leads to tissue growth. However unlike the previous formulation, we do not assume that the predominant growth process is crypt fission, but instead have individual crypts increase in size.

#### 2.1 Mathematical formulation

Let us consider an individual crypt and denote by $y_i$ the number of SCs in the crypt that have the different mutational status. Similar to the subscripts used in the previous model, $y_1$ corresponds to the wild-type cells, $y_2$ to the APC$^{+/-}$ cells, $y_3$ to the APC$^{-/-}$ cells, $y_4$ to the Kras$^-$ cells, $y_5$ to cells that have both APC$^{+/-}$ and Kras$^-$ status, and finally, $y_6$ to cells that comprise late adenoma and are characterized by both APC$^{-/-}$ and Kras$^-$ status. The model is presented schematically in figure 8, where the 6 types are denoted by circles, and their parameters are marked. In particular, we have the mutational processes identical to the ones described before. We also assume that cells of each type are characterized by a basic division rate and death rates, $r$ and $d$, and grow and compete according to a standard logistic process. We assume that different cell types can have different associated carrying capacity parameters, see figure 8: types $y_1$ and $y_2$ have the carrying capacity $K$; type $x_3$ has the carrying capacity $K_A$, where $K_A > K$; types $y_4$ and $y_5$ have the carrying capacity $K_R$, $K_R > K$. Type $y_2$ growth at the net rate $R$.

When formulating the logistic growth in an evolutionary system, one can make different assumptions. Healthy cells in a crypt are in a state of homeostasis, which is characterized by a relatively slow turnover, where self-renewing cell divisions are approximately balanced by differentiation divisions. In this model, self-renewal (differentiation) divisions mathematically act as divisions (deaths) in the birth-death process. The kinetic rates are modified by how close the population is to its homeostatic size (the carrying capacity), reflecting the effect of signaling, including cellular feedback control loops, that may orchestrate cell fate decisions within lineages. Two different modeling approaches were considered, which we refer to as "division-controlled growth" and "death-controlled growth", see [5].

##### 2.1.1 Division-controlled growth

In the division-controlled growth model we assume that as a cell population approaches its carrying capacity, it is the rate of self-renewal that decreases; at homeostasis, the division rate matches the death rate (this is what we termed "division-controlled growth" in [5]).

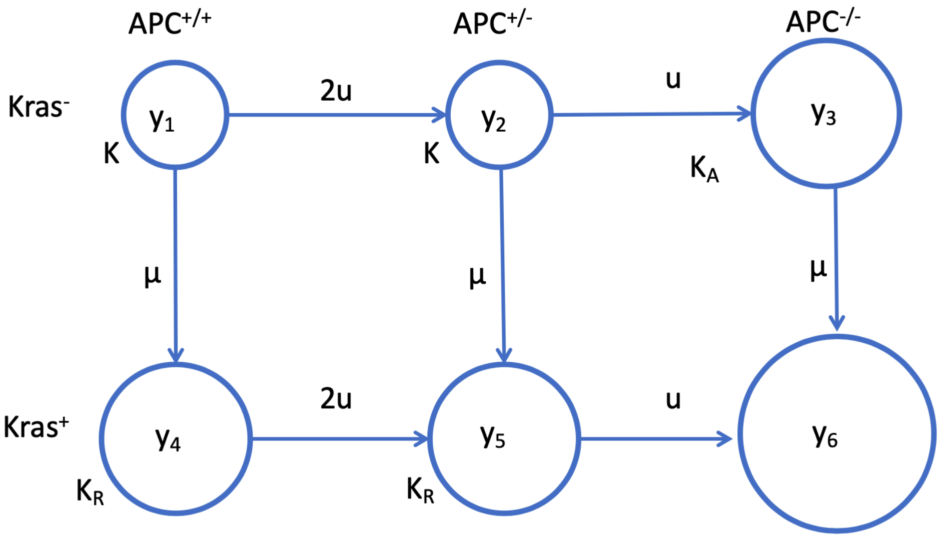

Figure 8: A schematic showing the relevant cell types, their fitness values, carrying capacities, and mutational processes that contribute to the generation of colon cancer.

Expressed as a deterministic system of ODEs, the dynamics can be written as follows:

$$
\begin{aligned}
\dot{y}_1 &= r y_1 (1-\mu-2u)\left(1-\frac{Z}{K}\right) - d y_1,\\
\dot{y}_2 &= 2ury_1\left(1-\frac{Z}{K}\right) + ry_2(1-\mu-u)\left(1-\frac{Z}{K}\right) - dy_2,\\
\dot{y}_3 &= ruy_2\left(1-\frac{Z}{K}\right) + r(1-\mu)\left(1-\frac{Z}{K_A}\right) y_3 - dy_3,\\
\dot{y}_4 &= r\mu y_1\left(1-\frac{Z}{K}\right) + r(1-2u)\left(1-\frac{Z}{K_R}\right) y_4 - dy_4,\\
\dot{y}_5 &= 2ur\left(1-\frac{Z}{K_R}\right) y_4 + r\mu\left(1-\frac{Z}{K}\right) y_2 + r(1-u)\left(1-\frac{Z}{K_R}\right) y_5 - dy_5,\\
\dot{y}_6 &= ru\left(1-\frac{Z}{K_R}\right) y_5 + r\mu\left(1-\frac{Z}{K_A}\right) y_3 + Ry_6,
\end{aligned}
\tag{26}
$$

where

$$Z = \sum_{i=1}^{5} y_i$$

In the absence of type $y_6$, the system grows to a steady state. If $y_6$ is created then growth becomes unrestricted.

This "division controlled growth" model is characterized by the following distinct properties. Suppose the resident population has parameters $r_1, d_1, K_1$. When the resident population size is much lower than its equilibrium size (given by $K_1(1-d_1/r_1)$), cells divide with rate $r_1$, which is greater than the death rate ($r_1 > d_1$), such that the population grows. As the population size approaches its equilibrium value, the division rate decreases until it reaches $d_1$. At steady state, the cells divide and die with rate $d_1$. This is in a sense similar to the constant-population Moran DB process (see e.g. [1]). A mutant cell that is characterized by kinetic parameters $r_2, d_2, K_2$, when it is first produced inside the resident population at equilibrium, has the initial growth rate $r_2\left(1-\frac{K_1}{K_2}\left(1-\frac{d_1}{r_1}\right)\right)$. The relative fitness of the mutant is given by

$$\frac{r_2}{d_1}\left(1-\frac{K_1}{K_2}\left(1-\frac{d_1}{r_1}\right)\right)\frac{d_1}{d_2}.$$

If we assume that $r_1 = r_2 = r$ and $d_1 = d_2 = d$, and the mutant fitness advantage is only given by a larger carrying capacity associated with the mutant, then the expression for the relative mutant fitness becomes

$$\frac{r}{d}\left(1-\frac{K_1}{K_2}\left(1-\frac{d}{r}\right)\right), \tag{27}$$

and if $K_2 \gg K_1$, it can be as large as $r/d$.

###### 2.1.2 Death-controlled growth

The death-controlled growth model assumes that as the population grows closer to the carrying capacity, it is the differentiation and/or death rate increases to match the self-renewal rate ("death-controlled growth" of [5]). This can be expressed as a system of ODEs in the following way:

$$\begin{aligned}
\dot{y}_1 &= ry_1(1-\mu-2u) - dy_1\left(1+\frac{Z}{K}\right),\\
\dot{y}_2 &= 2ury_1 + ry_2(1-u-\mu) - dy_2\left(1+\frac{Z}{K}\right),\\
\dot{y}_3 &= ruy_2 + r(1-\mu)y_3 - dy_3\left(1+\frac{Z}{K_A}\right),\\
\dot{y}_4 &= r\mu y_1 + r(1-2u)y_4 - dy_4\left(1+\frac{Z}{K_R}\right),\\
\dot{y}_5 &= 2ury_4 + r\mu y_2 + r(1-u)y_5 - dy_5\left(1+\frac{Z}{K_R}\right),\\
\dot{y}_6 &= ruy_5 + r\mu y_3 + Ry_6.
\end{aligned} \tag{28}$$

Note that the equilibrium value in the wild type compartment in division-controlled system (26) was $K(1-d/r)$, and the division rate of the $y_1$ cells at equilibrium was given by $d$. In death-controlled system (28), the equilibrium population is $K(r/d-1)$, and the equilibrium division rate is $r$. To have the same equilibrium and turnover rate in system (28) as in system (26), we would have to use the following replacement in system (28):

$$r \to d, \quad d \to \frac{d}{2-d/r}$$

The "death controlled growth" model is characterized by the following properties. Suppose as before the resident population has parameters $r_1, d_1, K_1$. When the resident population size is much lower than its equilibrium size (given by $K_1(r_1/d_1 - 1)$), cells divide with rate $r_1 > d_1$. As the population size approaches its equilibrium value, the death rate increases until it reaches $r_1$. At steady state, the cells divide and die with rate $r_1$, and this is similar to the constant-population Moran BD process (see e.g. [1]). A mutant cell that is characterized by kinetic parameters $r_2, d_2, K_2$, when it is first produced inside the resident population at equilibrium, has the growth rate of $r_2$ and the initial death rate $d_2\left(1 + \frac{K_1}{K_2}\left(\frac{r_1}{d_1} - 1\right)\right)$. The relative fitness of the mutant is given by

$$\frac{r_2}{d_1}\frac{d_1}{d_2\left(1 + \frac{K_1}{K_2}\left(\frac{r_1}{d_1} - 1\right)\right)}.$$

If we assume that $r_1 = r_2 = r$ and $d_1 = d_2 = d$, and the mutant fitness advantage is only given by a larger carrying capacity associated with the mutant, then the expression for the relative mutant fitness becomes

$$\frac{r}{d}\frac{1}{\left(1 + \frac{K_1}{K_2}\left(\frac{r}{d} - 1\right)\right)}, \tag{29}$$

and if $K_2 \gg K_1$, again it can be as large as $r/d$.

#### 2.2 Stochastic dynamics: Gillespie approach and the coarse-grained approximation

To incorporate stochasticity into the description, we used Gillespie simulations. Given the system parameters, many runs were performed to calculate the probability to create a mutant of type y6 by time $t$, $p(t)$. The main objective of the mathematical modeling of these processes is to study how the incidence curve depends on the different parameters, which requires a very large number of simulations. To cut simulation time, we used an approximation inspired by the methodology that we developed in [3, 4] in a different context and for a different underlying process, the constant-population Moran process. The idea is to represent the dynamics as a sequence of transition between a small number of "coarse-grained" states, which are characterized by fixation of different types. For example, we can denote by $X_1(t)$ the probability that the whole population consists of normal cells; $X_2(t)$ the probability that the whole population consists of cells with one copy of the AC gene inactivated; $X_4(t)$ the probability that the whole population consists of cells with KRAS activated (and wild type APC), etc. The simplifying assumption is that the system spends most of its time in one of these states, and intermediate states that contain heterogeneous population are short-lived. In terminology developed in [3, 4], transitions $X_1 \to X_2 \to X_3$ represent processes of sequential fixation of $APC^{+/-}$ and then $APC^{-/-}$ mutants, and transition $X_1 \to X_3$ represents stochastic tunneling whereby a successful mutant of type $APC^{-/-}$ is created from an $APC^{+/-}$ clone although the latter never reaches fixation. This approximation allows one to obtain function $p(t)$ by simply solving a system of 6 ordinary differential equations (which are different from either of the systems (26) or (28)), which leads to a very significant (orders of magnitude) cut in computational time. The validity of this methodology was established by repeated simulations.

We obtained a coarse-grained model to describe the evolutionary system based on the Moran process, see [3] In the Moran process, we assume the total population stays constant over time. At each time step, a random cell type is chosen for division proportional to fitness, and a random cell type is chosen for death. A mutation cell may be produced in the division event with a small probability.

We use $X_i(t)$, $1 \leq i \leq 6$ to denote the probabilities at time $t$ that the system consists cells exclusively of type $y_i$. The Kolmogorov forward equations for the system (26) are as follows:

$$\begin{aligned}
\frac{dX_1}{dt} &= -X_1(t)(R_{12} + R_{13} + R_{14} + R_{15}), \\
\frac{dX_2}{dt} &= X_1(t)R_{12} - X_2(t)(R_{23} + R_{25} + R_{26}), \\
\frac{dX_3}{dt} &= X_1(t)R_{13} + X_2(t)R_{23} - X_3(t)R_{36}, \\
\frac{dX_4}{dt} &= X_1(t)R_{14} - X_4(t)(R_{45} + R_{46}), \\
\frac{dX_5}{dt} &= X_1(t)R_{15} + X_2(t)R_{25} + X_4(t)R_{45} - X_5(t)R_{56}, \\
\frac{dX_6}{dt} &= X_2(t)R_{26} + X_3(t)R_{36} + X_4(t)R_{46} + X_5(t)R_{56}.
\end{aligned} \tag{30}$$

Here, similarly to the previous model, $R_{ij}$ stands for population transition rate from state $X_i$ to state $X_j$. The incidence curve (that is, the probability that a late adenoma occurred at time $t$) is then calculated as

$$p(t) = 1 - (1 - X_6)^{N_{crypt}}. \tag{31}$$

There are both similarities and differences between this approach and that of systems (1-5) and (19-23). The similarity is the underlying assumption that each crypt is with a high probability homogeneous with respect to mutations. This is the basis of both approaches. The differences are as follows:

- The approach of (1-5) and (19-23) studies populations of crypts, and that of system (30) studies a single crypt. Therefore, the variables in systems (1-5) and (19-23) refer to the numbers of crypts of different types ($n_i$), and in system (30) they are probabilities for a single crypt to be of a given type ($X_i$).
- Crypts in the approach of (1-5) and (19-23) are assumed to be of a constant size, but they can divide thus increasing their number; a fitness advantage conferred by mutations is expressed as an increase crypt fission rate. Crypts in the approach of system (30) can be of different sizes (but cannot divide); a fitness advantage conferred by mutations is expressed as an increase in a crypt size.

- The transition rates in the approach of (1-5) and (19-23) are calculated by using the methods of [3, 4]. As a consequence of crypt size differences in the approach of system (30), however, the methodology must be adjusted.

Below we describe the procedure of transition rate calculations for the division-controlled and death-controlled growth models. The main difference between the current case and the models of [2–4] is the fact that in the present system the population grows as a consequence of mutational fixation, and thus the constant-population Moran process cannot be used. Instead, in the simulations we used a Gillespie method based on logistic-growth ODEs. The transition rates for the coarse grained approach are calculated as described below.

##### 2.2.1 Coarse-grained approach for division-controlled model

Let us first suppose that we calculate a rate between compartments $x_{res}$ and $x_{inv}$, $R_{x_{res},x_{inv}}$, where the subscripts *res* and *inv* stand for "resident" and "invader", respectively. If a rate is a transition between neighboring compartments, that is, if type $j$ is produced from type $i$ as a result of a single mutational event, then the corresponding mechanism falls under "sequential fixation:" pattern, whereby a mutant of type $j$ (the "invader") is produced in the population of type $i$ (the "residents") at equilibrium, and with a certain probability the invader takes over (reaches fixation). In this case,

$$R_{x_{res},x_{inv}} = N_{res} r_{res} u_{res-inv} \rho_{inv,resi},$$

where $N_{res}$ is the steady state size of compartment $x_{res}$, $u_{res-inv}$ is the full mutation rate that creates type $x_{inv}$ from type $x_{res}$, and $\rho_{inv,res}$ is the probability that starting from 1 cell of type $x_{inv}$ among $N_{res}$ cells of type $x_{res}$, type $x_{inv}$ reaches fixation. This probability is given by

$$\rho_{inv,res} = \begin{cases} \frac{1-1/f_{inv,res}}{1-1/f_{inv,res}^{N_{inv}}}, & f_{inv,res} \neq 1, \\ \frac{1}{N_{inv}}, & f_{inv,res} = 1, \end{cases} \tag{32}$$

where $N_{inv}$ is the steady-state size of compartment $x_{inv}$ and $f_{inv,res}$ is a relative fitness of type $x_{inv}$ compared to type $x_{res}$. We have

$$f_{inv,res} = \frac{r_{inv} d_{res}}{d_{inv} r_{res}},$$

where $r_{res}, d_{res}$ are division and death rates of the resident type at equilibrium, and $r_{inv}, d_{inv}$ are division and death rates of invading type, when introduced at low numbers among the resident cells at equilibrium.

For example, to calculate $R_{x_1 x_2}$, we note that this is a sequential fixation rate, because type $x_2$ is produced from type $x_1$ with mutation rate $u_{res-inv} = u$. We further have $N_{res} = K\left(1 - \frac{d}{r}\right)$ and $N_{inv} = K_A\left(1 - \frac{d}{r}\right)$. To calculate the relative fitness of type $x_2$ compared to type $x_1$, we have to compare their kinetic rates. This can be done by considering a system of ODEs that describes the coexistence dynamics of the two types. We have (in the absence of mutation rates)

$$\dot{x}_1 = r x_1 \left(1 - \frac{x_1 + x_2}{K}\right) - d x_1, \quad \dot{x}_2 = r x_2 \left(1 - \frac{x_1 + x_2}{K_A}\right) - d x_2.$$

At the host equilibrium, $x_1 = K\left(1-\frac{d}{r}\right)$ and $x_2 = 0$, the host cells' divisions are balance with their deaths, such that

$$r_{res} = d, \quad d_{res} = d.$$

Note that the expression for the effective division rate of the host, $d_i$, can be obtained by using the steady state values of $x_1, x_2$ in the expression

$$r_{res} = r\left(1 - \frac{x_1 + x_2}{K}\right) = d.$$

For the invader, we assume that $x_2 \ll x_1$ and again use the equilibrium values to calculate

$$r_{inv} = r\left(1 - \frac{x_1 + x_2}{K_A}\right) = r\left(1 - \frac{K}{K_A}\left(1 - \frac{d}{r}\right)\right), \quad d_{inv} = d.$$

Therefore, we obtain

$$f_{inv,res} = \frac{r}{d}\left(1 - \frac{K}{K_A}\left(1 - \frac{d}{r}\right)\right). \tag{33}$$

We note that if $K_A > K$, $f_{inv,res} > 1$, that is, the invader is advantageous. The sequential fixation rate is then given by

$$R_{x_1 x_2} = K\left(1 - \frac{d}{r}\right) du \rho_{x_2,x_1},$$

with $\rho_{x_2,x_1}$ given by equations (32,33).

To calculate rate $R_{x_0,x_1}$, which is again a sequential fixation rate, we note that $u_{res-inv} = 2u$, and $N_{res} = N_{inv} = K\left(1-\frac{d}{r}\right)$. We further note that since the carrying capacity of the two compartments is the same, $f_{inv,res} = 1$, and $\rho_{inv,res} = 1/N_{inv}$. We therefore have a sipler formula,

$$R_{x_0 x_1} = 2du.$$

The second type of rates connects compartments that are not immediately adjacent and represents the process of stochastic tunneling. In this scenario, a mutant clone could be produced by a mutation in the resident compartment, but before the mutant clone reaches fixation, one of its cells experiences a second mutation, and the resulting double-hit mutant expands and reaches fixation, displacing the resident population. In our system, the tunneling rates are $R_{x_0 x_2}, R_{x_0 y_1}, R_{y_0 y_2}, R_{x_1 y_2}$. The tunneling rates can be calculated, and simple expressions exist for several limiting cases. In particular, the relative fitness of the intermediate type is important. If the intermediate type has the same fitness as the resident type, we have neutral tunneling; this is the case for rates $R_{x_0 x_2}, R_{x_0 y_1}$ (through $x_1$), $R_{y_0 y_2}$. In the case of neutral tunneling, the rate is given by

$$R_{x_{res} x_{inv}} = N_{res} r_{res} u_{res-int} \sqrt{u_{int-inv}} \rho_{res,int},$$

where subscript *int* refers to the intermediate phenotype.

For example, for the rate $R_{x_0 x_2}$, we have

$$R_{x_0 x_2} = K\left(1 - \frac{d}{r}\right) d\, 2u \sqrt{\mu} \rho_{x_2,x_0}.$$

##### 2.2.2 Coarse-grained approach for death-controlled model

There are several changes that enter the calculation of the coarse-grained rates in the death-controlled growth compared to the division-controlled growth.

- Population sizes are given by expressions of the type

$$N = K\left(\frac{r}{d} - 1\right).$$

- The division and death rates or resident and invader types change. We have

$$r_{res} = r, \quad d_{res} = r,$$

  because it is the death rate that adjusts to match the division rate near the carrying capacity. Further, we have for the invader type:

$$r_{inv} = r, \quad d_{inv} = d\left(1 + \frac{K_{res}}{K_{inv}}\left(\frac{r}{d} - 1\right)\right).$$

- As a consequence, the relatie fitness of the invader is given by

$$f_{inv,res} = \frac{r}{d}\left(1 + \frac{K_{res}}{K_{inv}}\left(\frac{r}{d} - 1\right)\right)^{-1}$$

##### 2.2.3 Comparison between the coarse-grained approximation and the Gillespie simulations

To measure the difference between the coarse-grained model and stochastic Gillespie simulations, we calculated the difference between the incidence curves obtained by the two methods, and defined the following relative $L_2$ error function:

$$Error = \frac{\sqrt{\sum_{k=1}^{n}(G_k - C_k)^2}}{\sqrt{\sum_{k=1}^{n}(G_k)^2}}, \tag{34}$$

where $k$ denotes the indices of all the selected discrete points along the incidence curves $G$ and $C$, obtained from the Gillespie and coarse-grained model respectively.

To test the applicability of the coarse-grained model, we construct a testing $2D$ parametric space that depends on the carrying capacity $K_A, K_R$ and the ratio between death and fitness rates $d/r$. Particularly, we set $K_A = K_R$ to vary in the set $\{101, 1001, 10001, 100001, 1000001\}$, and $d/r$ in the set $\{0.01, 0.1, 0.2, 0.3, 0.4, 0.5, 0.6, 0.7\}$, while fixing the initial population of $x_0$ and its carrying capacity $K$ to 100. The mutation rates of gene APC inactivation and the oncogene (Kras) are chosen to be physiological values, i.e. $10^{-7}$ and $10^{-9}$ respectively. We compared the incidence curves and report the $L_2$ error of the coarse-grained model for each parameter pairs in Figure 9 and 10.

To summarize, we obtained a good match between the coarse-grained and Gillespie approaches for both models, as long as the $d/r$ ratio was less than 0.5. For higher $d/r$ rations,

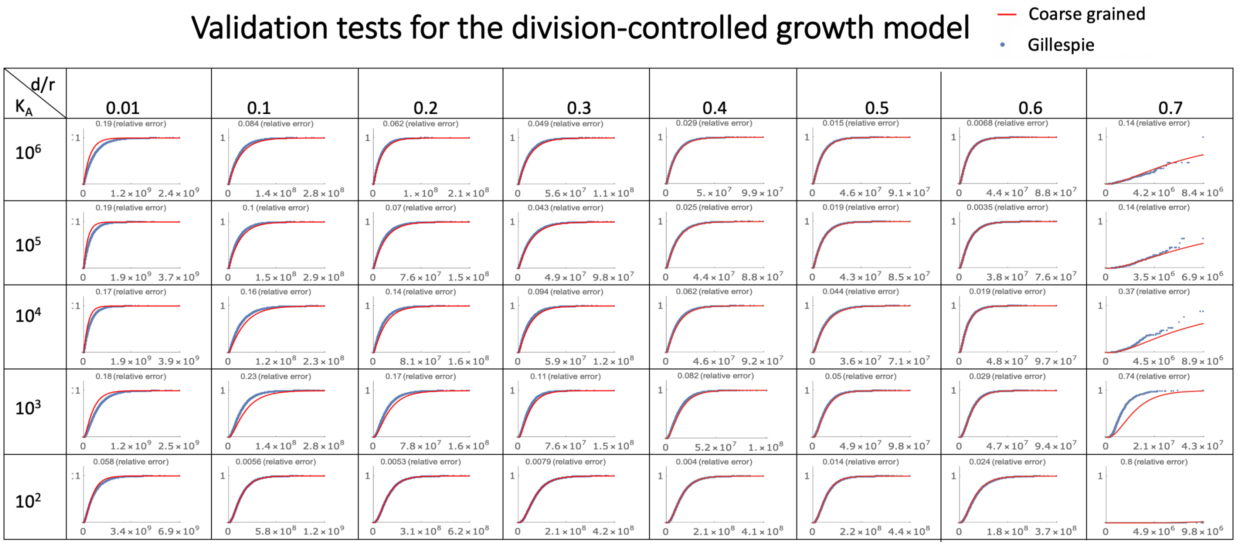

Figure 9: Comparison between the coarse-grained formulation and the Gillespie simulations for the division-controlled growth model. The carrying capacity $K_A = K_R$ and the ratio $d/r$ are varied. For each plot, 100,000 stochastic runs were performed. Nearly 100% simulations were successful for all cases, except when $d/r = 0.7$, where up to 90% of simulations showed extinction.

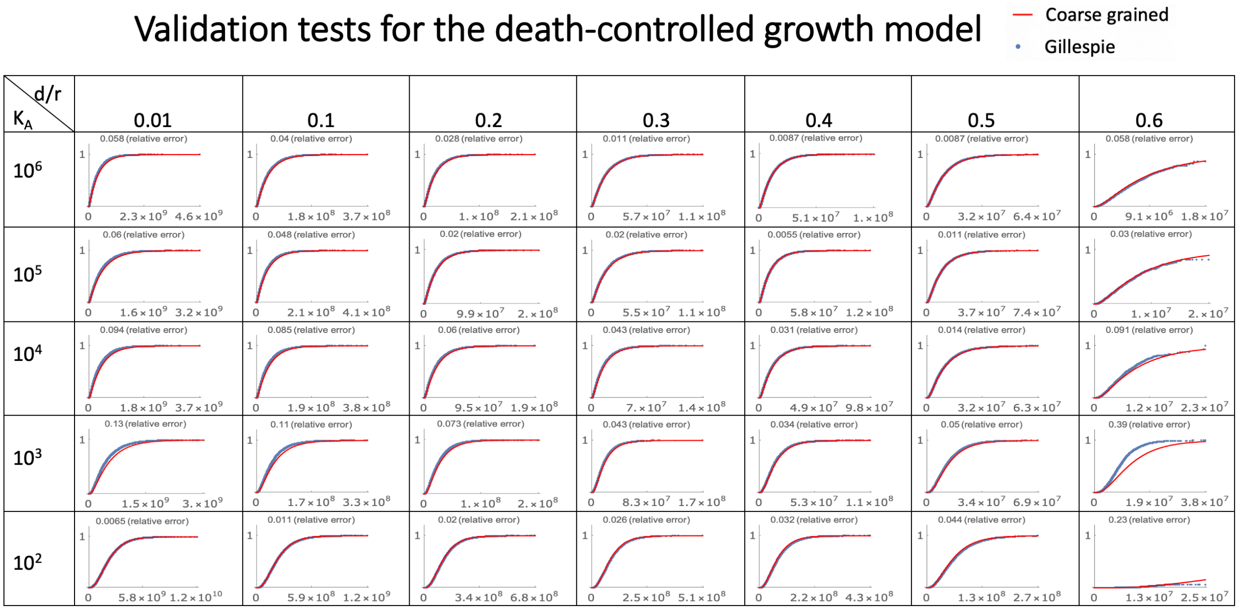

Figure 10: Same as figure 9 but for the death-controlled growth model. If $d/r = 0.6$, more than 90% of simulations showed extinction.

full extinction events in the Gillespie simulations become more common. Similarly, very small compartment size leads to extinction events in the Gillespie simulations.

In what follows we perform model fitting to the real-life age incidence curves for late adenoma. To do this, we use the coarse-grained approach because it provides an accurate description of the stochastic simulations in the parameter regions where Gillespie runs are stable, and it also allows to study the evolutionary dynamics outside those regions, including potentially biologically important regimes of small SC compartment size and relatively high turnover (larger $d/r$ ratios). In these regimes, the populations maintain stability due to control mechanisms that assure homeostasis, and whose details are not essential (apart from the fact that population survives and remains stable).

#### 2.3 Fitting the late adenoma data

The following procedure was used to fit the models to the late adenoma data. Parameters $u, \mu, N_{crypt}$ were fixed to their values in table 2, and the carrying capacity of the wild-type compartment, as well as the APC$^{+/-}$ compartment, $K$, was taken to be 10. The relative fitness of the most transformed type ($X_6$) with respect to the "host" type (the type where the last mutation happened) was taken 3.65 (which is the average between $F_3$ and $F_4$ in table 2). This value was used to calculate the rates $R_{i6}$ in system (30). The remaining parameters are $K_A, K_R, d$, and $r$. We however found it more convenient, instead of parameters $d$ and $r$, to work with parameters $d/r$ and $\tau$, where $\tau$ is the scaling of time. The carrying capacity parameters, $K_A$ and $K_R$, were each varied over 5 orders of magnitude. For each pair of $K_A$ and $K_R$, we varied parameter $d/r$ between 0 and 1 (with a step-size of 0.038), and for each $d/r$ found the best-fitting scaling parameter, $\tau$. We then accepted the resulting best fit if all of the following conditions were satisfied:

1. The resulting curve was within the values provided by males and females of the experimental cohort at age 73;
2. The cell division rate of the wild type at homeostasis was in the range of table 2;
3. The relative fitness of mutated cells was between 1.5 and 3.8.

Results of this fitting procedure are presented in figure 11 for the division-controlled growth and in figure 12 for death-controlled growth, respectively. Each panel corresponds to a specific combination of parameters $K_A$ and $K_R$ (indicated above each panel). The horizontal axis is time (yrs) and the vertical axis is the late adenoma probability. Blue and red dots represent the male and female data respectively (with the average given by purple dots). The lines (if any) correspond to the best fits selected as described above. We can see that satisfactory fits are available for a wide range of carrying capacity sizes, $K_A$ and $K_R$, and often include a range of $d/r$ values (several curves shown). Although the model is "easy" to fit (many good fits were found), the values of the carrying capacity parameters have to belong to a finite range (i.e. be above a threshold, and below another threshold), to ensure the realistic fitness advantage of mutants and to provide fits that are acceptable. In particular, very large carrying capacity values lead to an increase in the best fitting error. This is biologically similar to the conclusion from the previous model, where we saw that

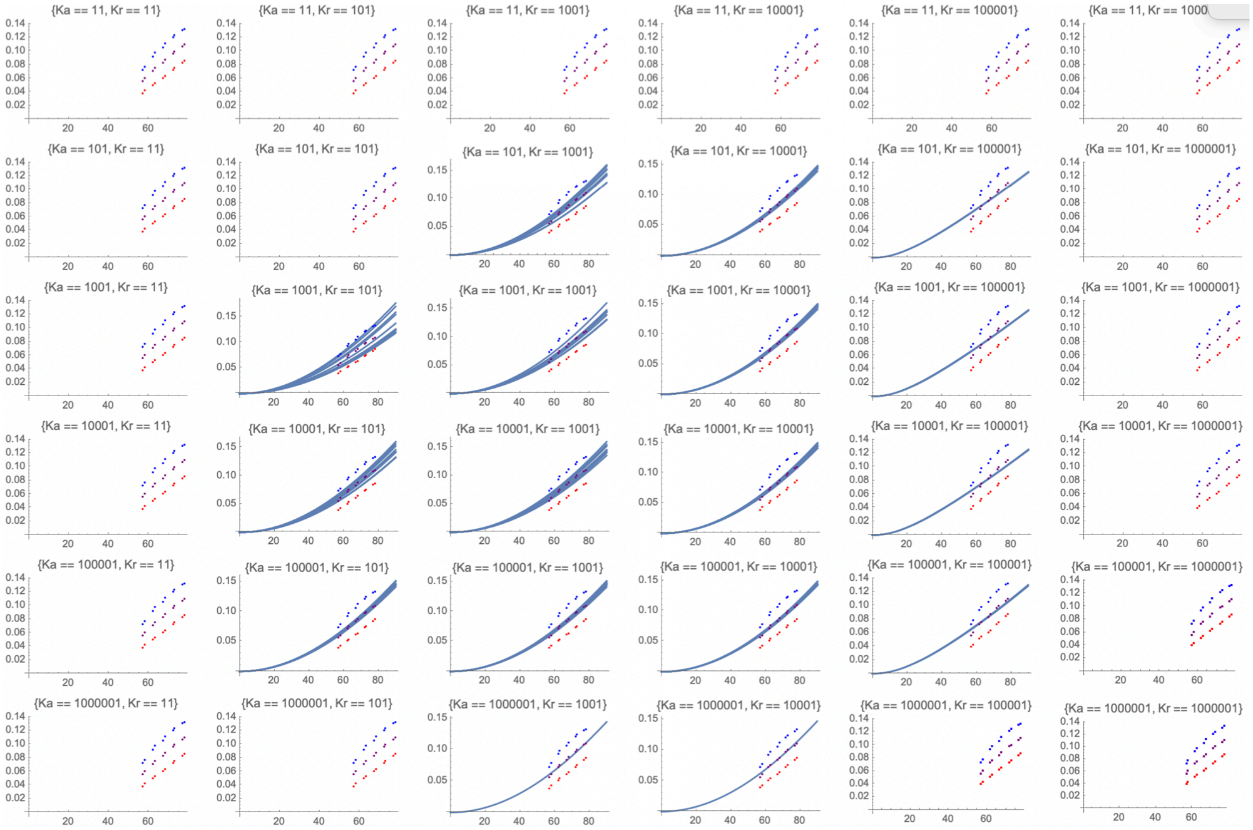

Figure 11: Best fits obtained by the division-controlled model. Each panel corresponds to a specific pair of parameters $K_A, K_R$ (indicated on top of each plot). For each plot, the horizontal axis is time (years) and the vertical axis is predicted (lines) and observed (dots) age-incidence of late adenoma (blue, red, and purple colors denote male, female, and mean). The different blue curves correspond to different $d/r$ values.

linear growth of crypts (without competition) could not result in an adequate fit to the late adenoma incidence curve.

Here we present predictions of the division-controlled and death-controlled models in the context of late adenoma formation and the delaying effect of aspirin.

Figure 13 shows the predicted fraction of late-adenomas that are created through the APC pathway, as obtained by the division-controlled and death-controlled models. The fractions are shown as heat-plots with darker colors corresponding to a higher prevalence of the APC pathway. The axes are the logarithms of the carrying capacity parameters. We can see that according to both models, a larger value of $K_A$ leads to a higher prevalence of the APC pathway. These results are similar to figure 3(d) of the main text, which was obtained by using model (19).

Figure 14 shows the reduction in the late-adenoma risk as a result of aspirin, as predicted by the two variants of the model. We explored two scenarios: (1) aspirin only affects the parameters of type 6 cells (blue dots) and (2) aspirin affects the modified types 3,4,5, and 6 (red dots). To implement the dose-dependent effect of aspirin, we varied parameters $r$ and $d$

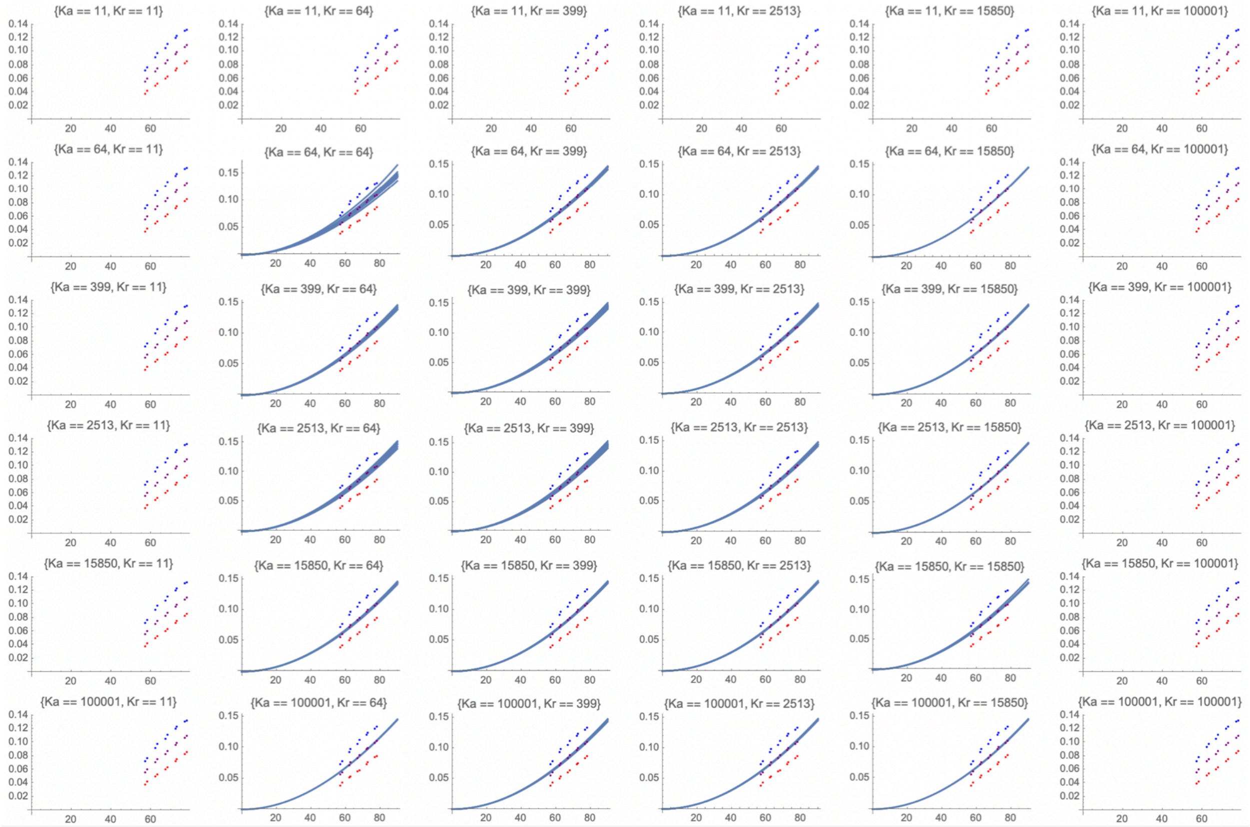

Figure 12: Same as in figure 11 but for the death-controlled model.

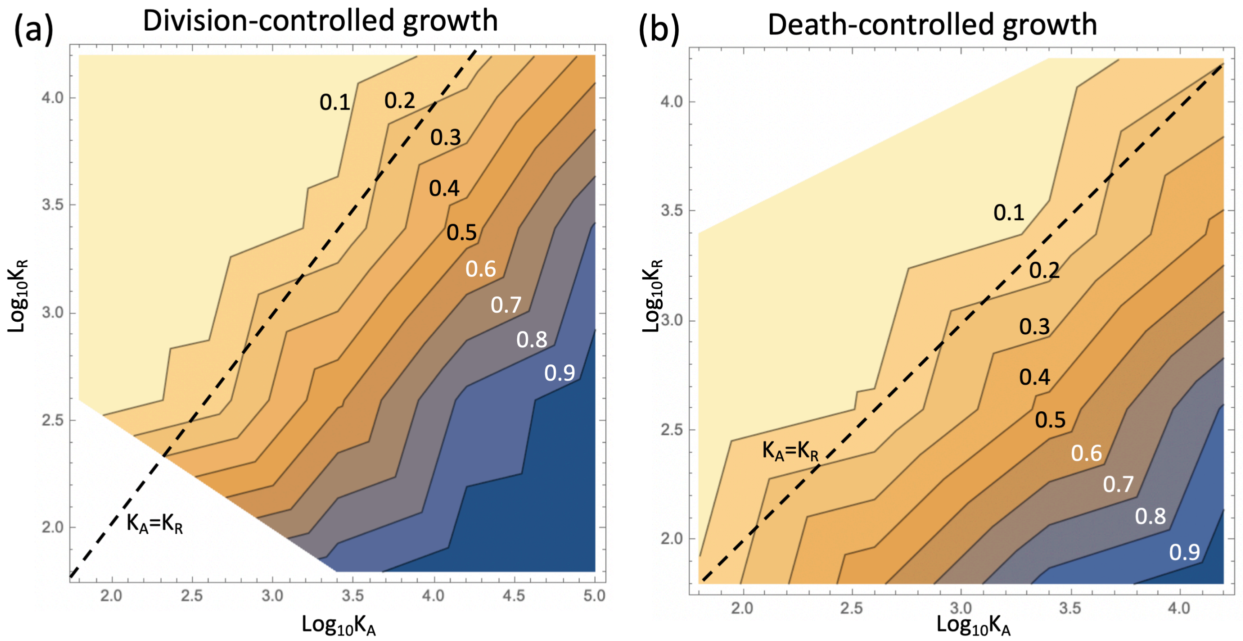

Figure 13: The proportion of late-adenoma incidence predicted to go through the APC-pathway. Darker colors indicate a larger proportion of the APC-pathway (and the fraction of the APC-path is marked by the contour lines). (a) Division-controlled model, (b) Death-controlled model. The dashed lines correspond to the two carrying capacity parameters equal to each other ($K_A = K_R$).

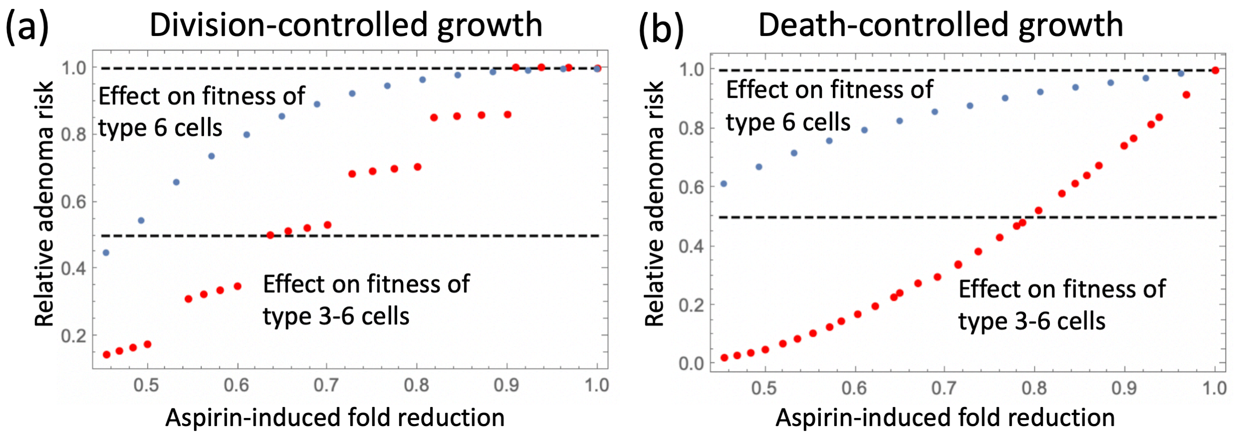

Figure 14: Relative adenoma risk under aspirin treatment, as a function pf aspirin-induced fitness reduction. (a) Division-controlled model, (b) Death-controlled model. Blue dots correspond to the model where only type 6 is affected by aspirin, and red dots to the model where types 3,4,5,6 are affected by aspirin.

of the affected cell types within the physiological limits, as discussed in the main text. The horizontal axis in figure 14 is a composite parameter that includes changes (if any) of the $r$ and $d$ values. Similar to figure 4 of the main text (which was obtained by using model (19)), we observe that for both models, biologically-relevant aspirin-induced fold reduction (between 0.45 and 1) leads to a decrease in the adenoma risk that is similar to that observed in the literature, if we assume that only type 6 cells are affected. Namely, if aspirin-induced fold reduction is around 0.5 then the risk of adenoma is about 50% reduced, which is the maximum observed effect. Again, similar to the observations of figure 4 of the main text, if aspirin affects other cell types, late adenoma risk reduction becomes stronger.
